# The genetic architecture of ecotypic differentiation in Chinook salmon of the California Central Valley

**DOI:** 10.1101/2025.07.26.666848

**Authors:** Eric C. Anderson, Neil F. Thompson, Anthony J. Clemento, Cassie Columbus, Ellen Campbell, Anne K. Beulke, John Carlos Garza

## Abstract

Understanding the genomic details underlying complex behavioral traits is a foundational pursuit in biology. We use genomic and genetic techniques to dissect the heritable underpinnings of adult migration timing of Chinook salmon in the California Central Valley (CCV), home to several ecotypes not found elsewhere. We find that a previously described genomic region contributes to the seasonal shift in adult freshwater migration in the CCV, as in other river basins, but we further identify two functional domains in this locus that separately and additively influence the trait, with each allele copy affecting timing by ∼two weeks. We show how the evolution of a unique ecotype in the CCV is partially due to an allele derived from the more widespread early-migrating haplotype. However, the genomic background of the evolutionarily differentiated ecotypes contributes a similar amount to trait variation. We show how a relatively simple five-allele genetic system, in concert with genomic backgrounds, can create a remarkable diversity of phenotypes and ecotypes for this iconic species.

**Teaser:** A diverse complex of salmon ecotypes is explained by a simple two-locus system and evolutionarily derived genomic backgrounds.

## Introduction

Many animals migrate to take advantage of different environments. Migration behavior involves an entire “syndrome” of traits necessary for its successful execution, and was thus traditionally viewed as being genetically complex (*1*). Prior to the advent of large-scale genome sequencing, studies of migratory behavior relied on quantitative genetic frameworks to resolve the degree of genetic control of migration (*2*), versus environmental factors, implicitly assuming a polygenic trait model. At the time, efforts to dissect the heritable component of migratory traits were largely confined to experimental manipulations on captive organisms, rather than observations of animals migrating through natural ecosystems (*3*). However, recent studies leveraging high-throughput sequencing have allowed genome-wide association studies of migratory behavior in natural populations, identifying supergenes and large-effect loci that influence migration propensity (*4*, *5*), timing (*6–8*) or direction (*9*) in fish and bird species.

Anadromous fishes, such as sturgeon, herring and salmon, are the archetypal migratory fishes, as their life history requires migration between freshwater and marine ecosystems (*10*). In two species of Pacific salmon (genus *Oncorhynchus*), a locus of major effect has been identified that is strongly associated with adult migration timing (*6–8*). This locus is in the vicinity of two genes, *GREB1L* and *ROCK1*, and is termed the *GREB1L/ROCK1* region (GRR). In Chinook salmon (*O. tshawytscha*), previous work found the part of this locus that is most strongly associated with categorical trait variation and termed it the Region of Strongest Association (RoSA), which carries haplotypes belonging to two broad groups associated with early or late migration in at least two lineages of the species (*8*). That work also identified a structural variant contiguous with the RoSA, but did not examine its association with trait variation. While there has been a focus on these “early” and “late” allelic lineages, and their effects on trait variation, the variation within the lineages has only recently begun to be explored (*11*). Little is known about the relative contributions of the RoSA subdomains on trait variation, and whether the causative polymorphisms are non-synonymous substitutions in the exonic regions or exercising their effects through non-coding regions, such as transcription factors in the intergenic region. Furthermore, although studies have shown that spawn timing can be highly heritable in salmonids, even in the absence of variation in the GRR (*12*, *13*), the relative effect of the RoSA on migration timing, compared to that of other variation in the genome, remains unknown.

The California Central Valley (CCV) is the southern natural range limit of Chinook salmon and harbors the greatest diversity of ecotypic variation in the species of any basin worldwide. Historically, CCV salmon migrated back to freshwater year-round, with distinct temporal modes that were categorized into ecotypes, or “runs” (*14*). Fall-run fish, currently the most numerous, have a modal return to their natal streams in October and November and spawn shortly thereafter. Spring-run fish enter freshwater primarily in the spring, remain in deep, cold-water pools over the summer, and proceed upriver to spawn in the fall, just prior to the mode of the fall-run spawning season. Historically, this life history gave the spring-run fish access to spawning grounds not easily reached by the fall-run fish, conferring a degree of spatial segregation and, in the CCV, genetic differentiation, from the fall-run ecotype that persists today (*15*). The winter-run ecotype arrives to freshwater in the late winter/early spring and spawns in the late spring and the early summer. It is highly genetically differentiated from all the other ecotypes, partially due to recent and extreme bottlenecks (*16*). Finally, late-fall-run fish begin arriving in late December and continue through February, spawning shortly after arrival. In spite of this notably distinct migration pattern, late-fall fish are only slightly genetically differentiated from the fall run (*15*).

Previous genome sequencing (*8*) showed that CCV fall-run and late-fall-run fish carry a late (*L*) lineage haplotype at the RoSA. Additionally, that work identified two distinct haplogroups amongst the early-lineage haplotypes, which we term here *E1* and *E2*. The *E1* haplotype was present throughout California spring-run populations, whereas *E2* was present only in the CCV, with the highest frequencies of it seen in the winter-run population. Importantly, this work also found that duplications in the intergenic region between *GREB1L* and *ROCK1* were highly associated with the haplotype lineages, but with a distinct pattern from the RoSA; the *L* and *E2* lineages were highly associated with the duplicated variant (*d*), whereas the *E1* lineage was almost always linked to the non-duplicated haplotype (*s*). However, the individual effects on adult migration timing of these different allelic lineages and of the intergenic duplications domain were unknown.

Water development and fishery management practices in the CCV have likely increased rates of interbreeding of early-and late-migrating fish relative to historical rates. At the large Feather River Hatchery (FRH) facility, it has been previously noted that, apart from the GRR, the spring-run fish and fall-run fish produced there are very similar genetically (*17*). This is likely a consequence of a long period of active selection for early-migrating fish, regardless of their lineage, for use in the reestablishment of a spring-run stock that was extirpated following dam construction. While naturally spawning spring-run fish, apart from those descended from FRH fish, can still be readily distinguished from the fall run using genetic markers (*15*), FRH Chinook salmon carry the early-lineage GRR alleles characteristic of early-migrating fish on a fall-run genomic background (GB).

Here, we directly estimate the effects of the different GRR lineages, the intergenic duplication alleles and the GB on adult migration timing of Chinook salmon in the CCV, providing a clear picture of the genomic architecture of this key phenotypic trait. This is made possible by the presence of multiple combinations of GRR alleles found in appreciable numbers in the CCV across all three GBs (winter, spring, and fall/late-fall), which provides a unique opportunity to disentangle the effects at the GRR from those of the broader genome.

To assess the genomic architecture of adult migration timing, we used genotype data for 5,651 adult Chinook salmon sampled on the date of arrival across 10 years at the Keswick Fish Trap (KFT), at the terminus of anadromy in the Sacramento River basin (Fig. 1). We used genetic stock identification to characterize the GB of each fish. Then, with newly developed genetic assays we determined the haplogroup allele (*E1*, *E2*, *L*) and duplication (*d*, *s*) genotypes for each fish and relate the genotypes to adult migration timing.

**Fig. 1:**
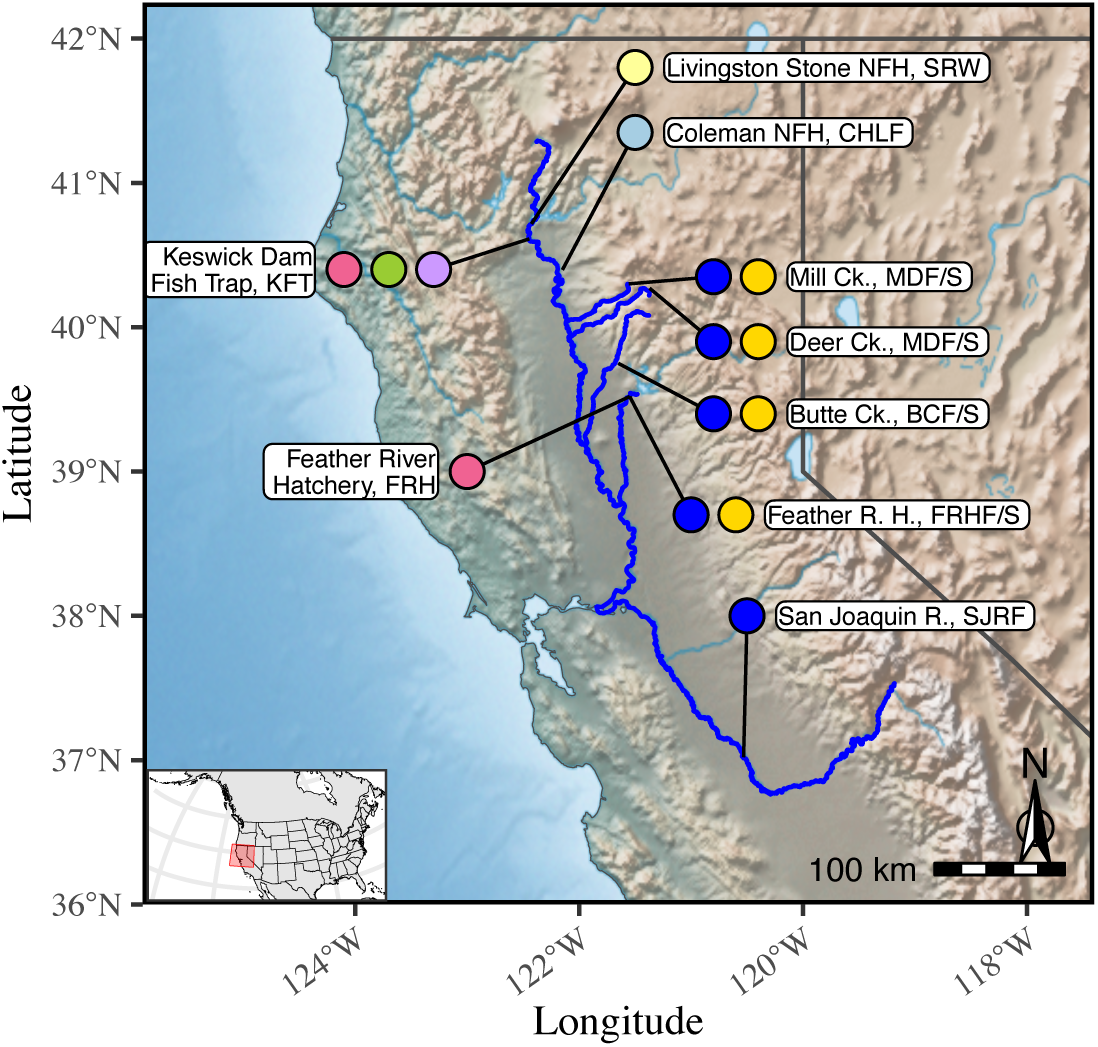
Map of sampling locations. Labels on the east side of the map represent populations used in reference baseline for assignment to genetic groups. Colored circles aside these location names indicate the different run-timing groups collected from each location: 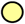 = Winter run, 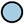 = Late-fall run, 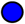 = Fall run, 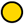 = Spring run. Names of each location are followed by the location codes. Labels on the west side of the map represent locations where samples of adult, migrating Chinook salmon were collected, with colored circles indicating the different genomic backgrounds (GB) of fish encountered at the site: 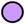 = Winter GB, 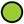 = Spring GB, 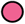 = Fall/late-fall GB. Abbreviations used in location names as follows: R. = River; Ck. = Creek; H. = Hatchery; NFH = National Fish Hatchery.

Our results indicate that both haplogroup allele (*E1*, *E2*, or *L*) as well as the presence or absence of the intergenic duplications (*d*, *s*), additively affect adult migration timing. These alleles combine into multiple genotypes with expected migration-timing effects that span a range of 100 days. These additive effects were corroborated in a second data set of 5,721 adult salmon arriving at the FRH, closer to the ocean in the same (Sacramento) river basin. We separately estimated the effect of the GB and evaluated candidate loci with large frequency differences between ecotypes for effects on the migration phenotype. Strikingly, we found the effect of GB to be nearly as large as the total additive effect of GRR alleles, but with no additional large-effect loci contributing to it, suggesting a substantial role for polygenic variation in Chinook salmon adult migration timing.

## Results and Discussion

We show here that the genomic architecture of the most diverse complex of Chinook salmon ecotypes in the species’ range is multifactorial with a remarkable contribution by a simple five-allele system in two tightly linked domains, within a locus of major effect, and with a significant contribution of genomic background (GB) associated with the three main evolutionary lineages in the basin. This major-effect locus, on chromosome Ots28, has been previously described as shifting seasonal migration timing of adult salmon in two species by several months. We show how two functional domains in this locus contribute additive effects to the migration timing trait in this species and how the evolution of a distinct sub-lineage in the early-migrating haplogroup has further diversified the temporal distribution of this critical phenotypic trait in one of the world’s iconic fish species.

Previous work has indicated that there are three main genetic lineages in the California Central Valley (CCV) Chinook salmon complex (Fig. 1): winter run, represented by a single remnant population; spring run, represented by several populations (Butte, Mill and Deer creeks) and fall run/late-fall run, which include the majority of salmon in the basin, including the phenotypically distinct late-fall run and the spring run in the Feather River subbasin (*17*). We confirmed the occurrence of these three lineages in our sample by comparison of values of *F*_ST_ between sampled populations, calculated both from amplicon-sequenced markers and whole-genome sequencing (Fig. 2A). We also confirmed that this 125-locus genotyping panel captures the major features of population structure that were resolved and identified with a larger panel that was recently described (*15*) (Fig. 2B). These features include that 1) winter run is substantially diverged from all other lineages; 2) the spring-run fish in Butte, Mill, and Deer creeks can be reliably distinguished from all fish in the basin; and 3) the spring-run fish at the Feather River Hatchery (FRH) are similar genetically to the fall-run and late-fall-run fish throughout the basin, likely due to a history of extensive cross-breeding between fall- and spring-run fish in the Feather River basin.

**Fig. 2:**
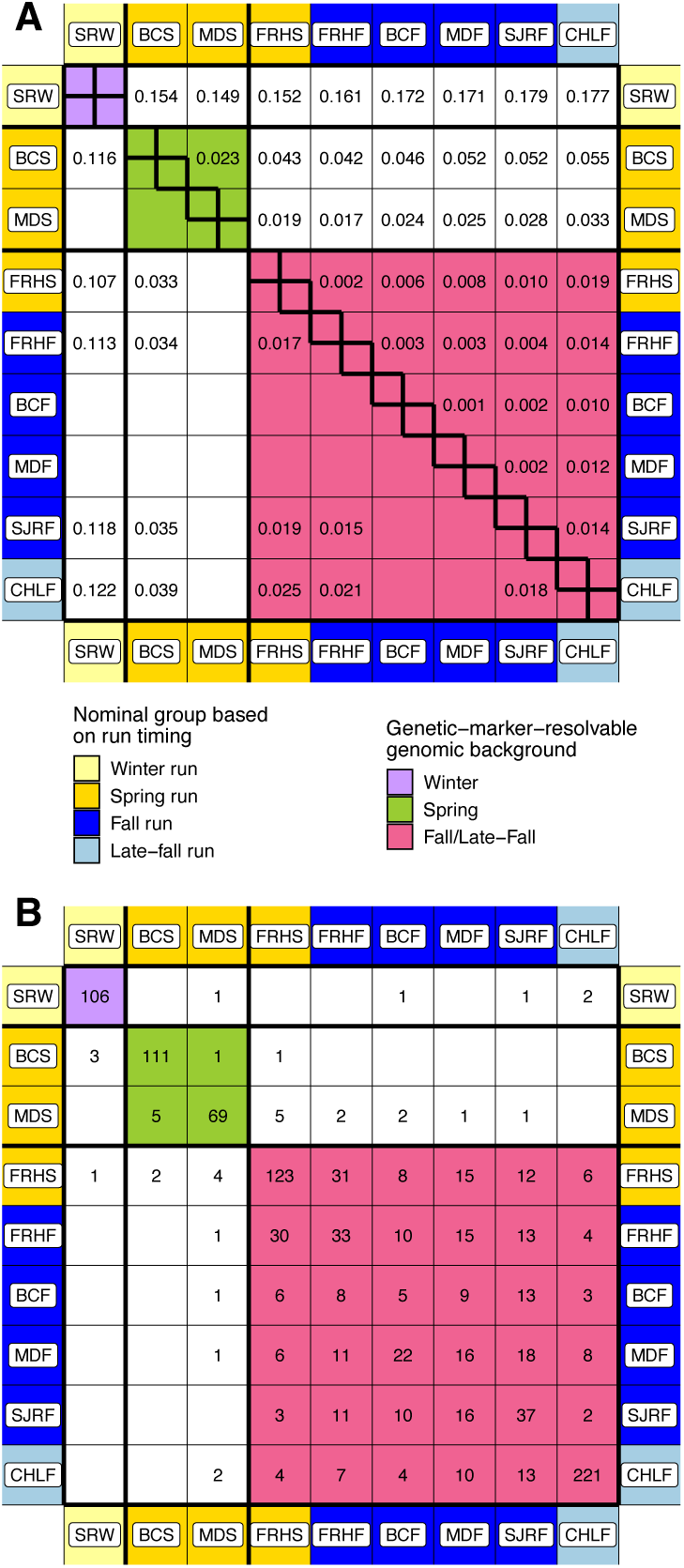
Pairwise. *F*_ST_ **values and genetic assignment rates.** Labels for populations of Chinook salmon in the California Central Valley are listed in Fig. 1. Colors on the perimeter give the nominal ecotype of each population based on run timing: yellow = winter run, gold = spring run, blue = fall run, light-blue = late-fall run. The colors in the interior highlight assignments between populations belonging to the three genomic backgrounds that can be resolved with these genetic markers: Purple = winter, green = spring; red = fall/late-fall. (**A**) Pairwise *F*_ST_ values between the populations. Above the diagonal are values calculated from the 125 genetic markers used for population assignment. Below the diagonal are *F*_ST_ values calculated from the whole-genome sequencing data available for only a subset of the populations. (**B**) Summary of population self-assignment results for a reference baseline of 1,073 typed at 125 genetic markers. Cells count the number of fish from a certain source population, denoted by the row labels, assigned to each population denoted by the column labels.

Genotyping 11,372 Chinook salmon samples (Tab. S1) from the Keswick Fish Trap (KFT) and the FRH’s spring-run program at 12 SNPs in the RoSA spanning 43.3 Kb (Tab. S2) provides an illuminating view of haplotype variation in this region. The 12 SNPs across this region were clearly organized into five haplotype groups (Fig. 3). Of the 11,198 fish with no missing data, all but 11 (<0.01%) carried a genotype composed of only these five haplotypes. The 11 that did not were likely due to genotyping errors or novel mutations, and were excluded from further analyses. The first 11 SNPs identify the *E1*, *E2*, and *L* alleles in the *E1E2L* domain (Fig. S1), within which we find no evidence for recombination. In contrast, we demonstrate that the distal SNP, RA7.T (Fig. 3), tags the intergenic duplications in the *sd* domain (Fig. S2) and our data provide clear evidence for recombination between the *E1E2L* and *sd* domains. Because of this recombination, haplotypes cannot be accurately resolved across both the *E1E2L* and *sd* domains in all individuals (Fig. 3). However, the variety of genotype combinations produced by recombination allows estimation of additive effects separately at the *E1E2L* and the *sd* domains. We note, however, that there is a complete lack of recombinant haplotypes that unite the *s* allele with the *L* allele in the two domains, in spite of many instances of heterozygotes where such recombination could occur. This raises the question of whether there is some mechanism, such as recombination suppression or selection on such recombinants, that is preventing their persistence in these populations. Also of note is that the *sd* domain has a distinct pattern of distribution across lineages relative to the rest of the genome in this group of fishes, with the *d* allele predominant in both the fall-run and winter-run lineages and the *s* allele present primarily in the spring-run lineage. In contrast, the rest of the genome is more similar between the spring-run and winter-run lineages, further suggesting that there may be a genomic incompatibility or other selective force acting on recombinants.

**Fig. 3:**
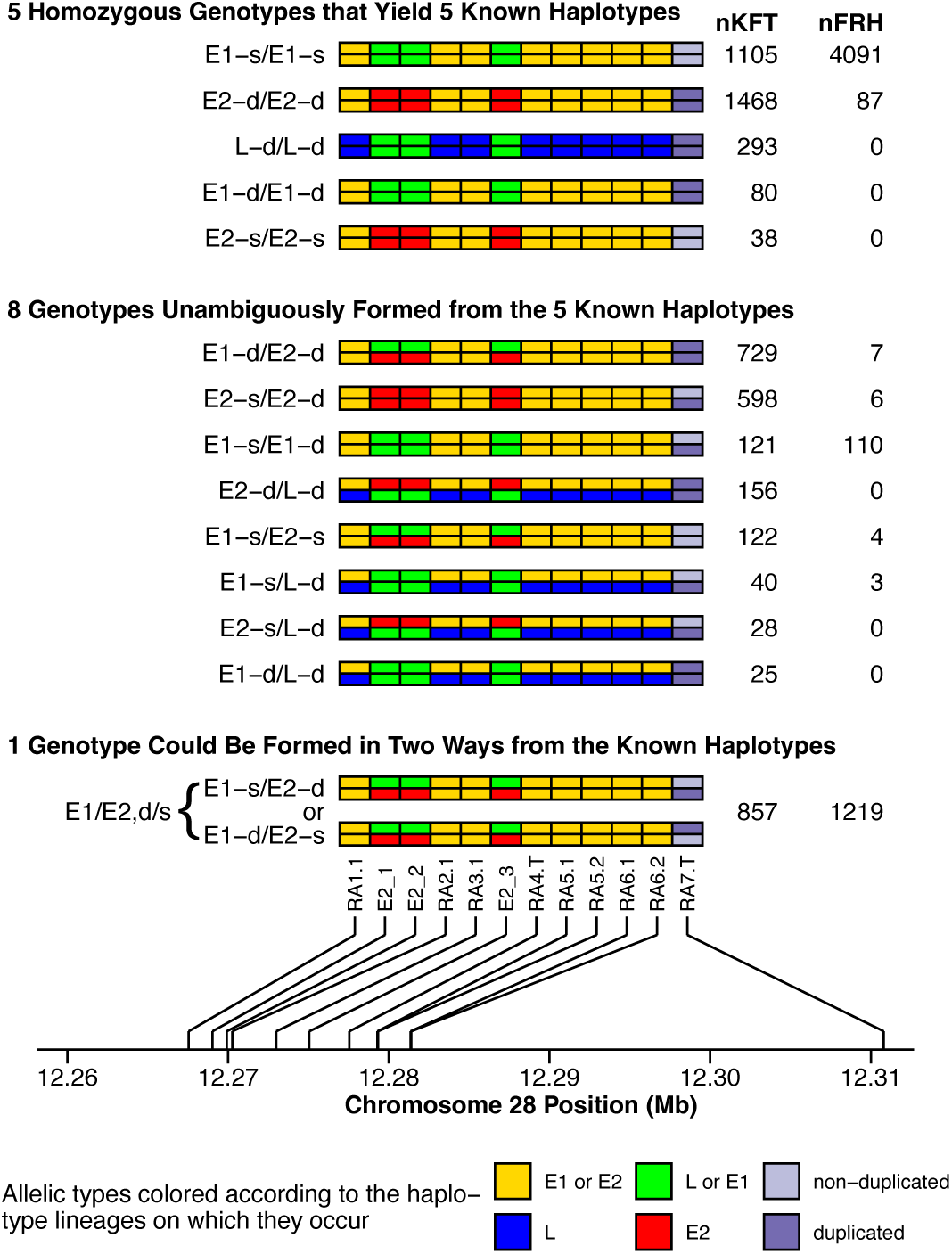
Resolvable haplotypes within the *GREB1L*/*ROCK1* region. There are three alleles, *E1*, *E2*, and *L* at the *LE1E2* domain, and two alleles, the duplicated (*d*) and non-duplicated (*s*) alleles at the *sd* domain. Each genotype of the 12 markers is shown as the combination of two haplotypes, one atop the other, the alleles within which are represented by colors within rectangular cells. The names and relative position of each variant are denoted by text names at the bottom and the lines connect those names to the Chromosome 28 position scale (in megabases). Names of the genotypes are given to the left of the genotype rectangles by the names of the haplotypes separated by a forward slash. For example, *E1-s/E1-s* is a homozygous genotype composed of two copies of the *E1-s* haplotype. Numbers to the right of the genotype rectangles show the number of such genotypes found at Keswick Fish Trap (nKFT), and at Feather River Hatchery (nFRH). Only one of the genotypes (*E1/E2,d/s*) cannot be resolved unambiguously into two haplotypes, and that is reflected in its name. We analyze the effects of this genetic variation in terms of the effects of each separate domain—one with alleles *E1*, *E2*, and *L*, and the other with alleles *d* and *s*.

A majority (4,178) of the 5,651 Chinook salmon sampled at KFT were genetically identified as carrying the winter GB, and these fish carried 14 different genotypes at the *E1E2L* and *sd* domains. The median migration times of the 14 different GRR genotypes on the winter GB varied by more than three months (Fig. 4), indicating a substantial contribution from the RoSA and duplication genotypes to adult migration timing. Inspection of the distribution of sampling dates across the KFT sampling window, from December 26 to August 1, in most years, indicates that the sampling window captures the vast majority of fish carrying the winter GB (Fig. S3).

**Fig. 4:**
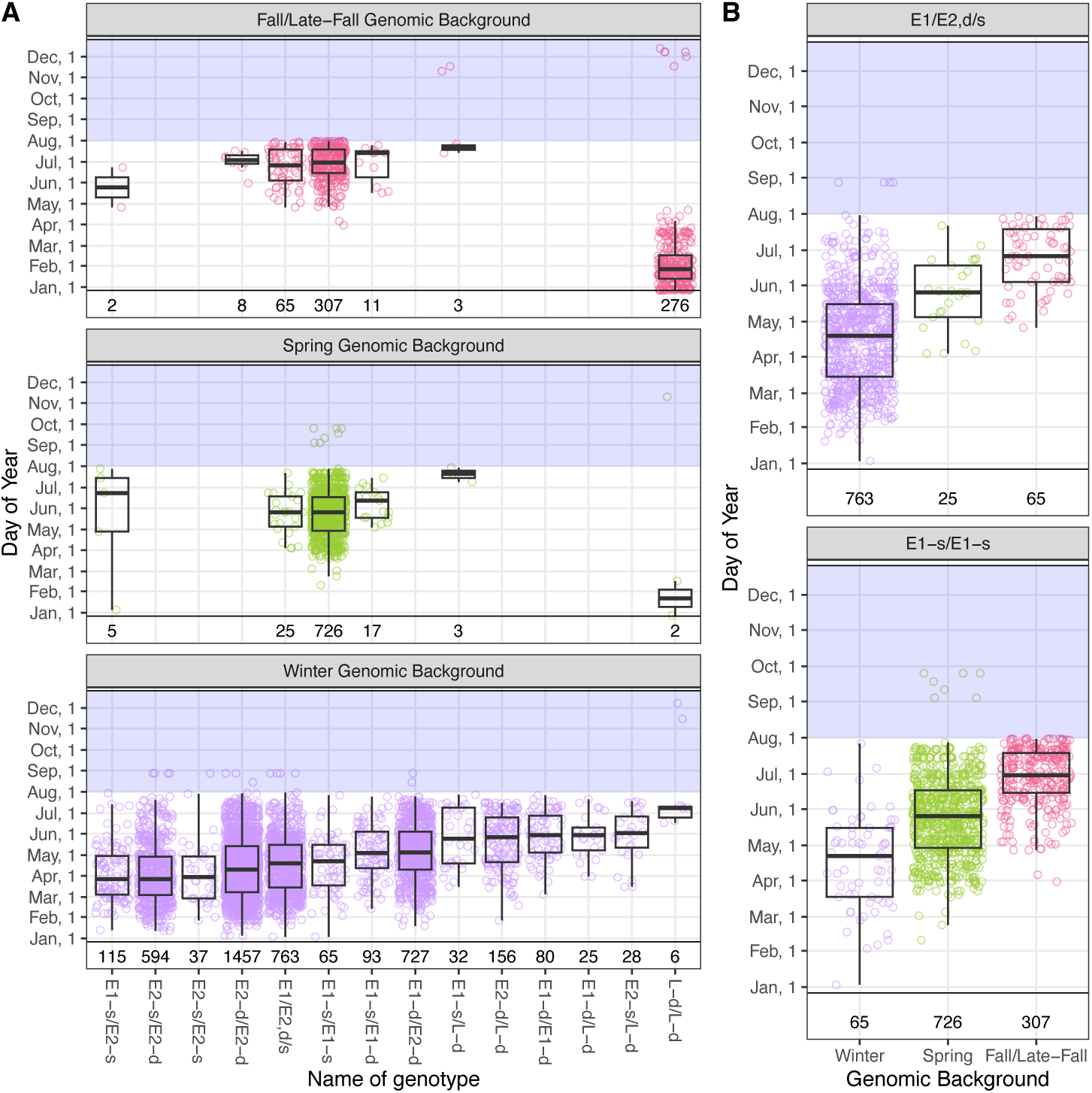
Arrival times at Keswick Fish Trap. (**A**) Along the *x*-axis are different genotypes at the two domains in the *GREB1L*/*ROCK1* region. The *y*-axis shows date of capture and sampling at the Keswick Fish Trap. Colors denote the different genetically discernible backgrounds: (Fall/Late-Fall, Spring, and Winter) as indicated in the facet headers. Sample sizes (number of genotyped fish) of each genotype appear at the bottom of each panel. The 276 fish on the right side of the Fall/Late-Fall panel are late-fall-run fish which were excluded from the analyses of the *GREB1L*/*ROCK1* region’s effect on migration timing. (**B**) Arrival times for fish of two different genotypes—*E1/E2,d/s* and *E1-s/E1-s*—for which sufficient samples were available for analysis of the effect of genomic background (GB) on migration timing. The three GBs are arranged along the x-axis, migration time is shown on the y-axis. The transparent blue rectangle represents the period when the trap does not typically operate. Values within that rectangle represent odd sampling dates within just one or two years and are discarded from analysis and in calculation of the boxplots. Dark lines are the median of sample points, the hinges are the first and third quartiles and the whiskers extend to the lowest and highest points less than 1.5 times the interquartile range from the lower or upper hinge, respectively. All sample points are plotted under each boxplot.

We treated sampling year as a random effect, and modeled the effect of the *E1*, *E2*, *L*, *s*, and *d* alleles within the winter GB at KFT, using a quantitative genetic framework suitable for multiple alleles. This revealed significant additive genetic effects of the *E1E2L* and *sd* domains on adult migration timing, with no significant dominance effects (Tab. S3). At the *E1E2L* domain, we chose the *E2* allele, which is predominant in winter-run fish, as the reference. The additive effect of each *E1* allele, relative to the reference, is an adult migration time 16.7 days later, while the additive effect of each *L* allele is 32.5 days. At the *sd* domain, the *s* (no duplications) allele was chosen as the reference, and the additive effect of each *d* allele (which carries the intergenic duplications) is adult migration timing 12.8 days later (Fig. 5A). Very similar results were obtained when using an animal model (*18*, *19*) by including the breeding values of all individuals as random effects in the model and supplying the genomic relationship matrix (GRM) calculated from the 125 marker loci to parameterize the variance/covariance structure of the breeding values. While the GRM cannot be estimated with great precision from such a small number of markers, this does still indicate that our estimated effects are not an artifact of unmodeled close relationships amongst our samples (Tab. S4; Fig. S4). Including the GRM also allows us to obtain an estimate of 0.034 for the narrow-sense heritability of the polygenic component for adult migration timing. Although this estimate is likely biased downward by the effect of unidentified relatives (*20*), it suggests that, within the winter GB, the large majority of variation in adult migration timing is additive and attributable to this region of chromosome Ots28, with little additional polygenic variation affecting this trait.

**Fig. 5:**
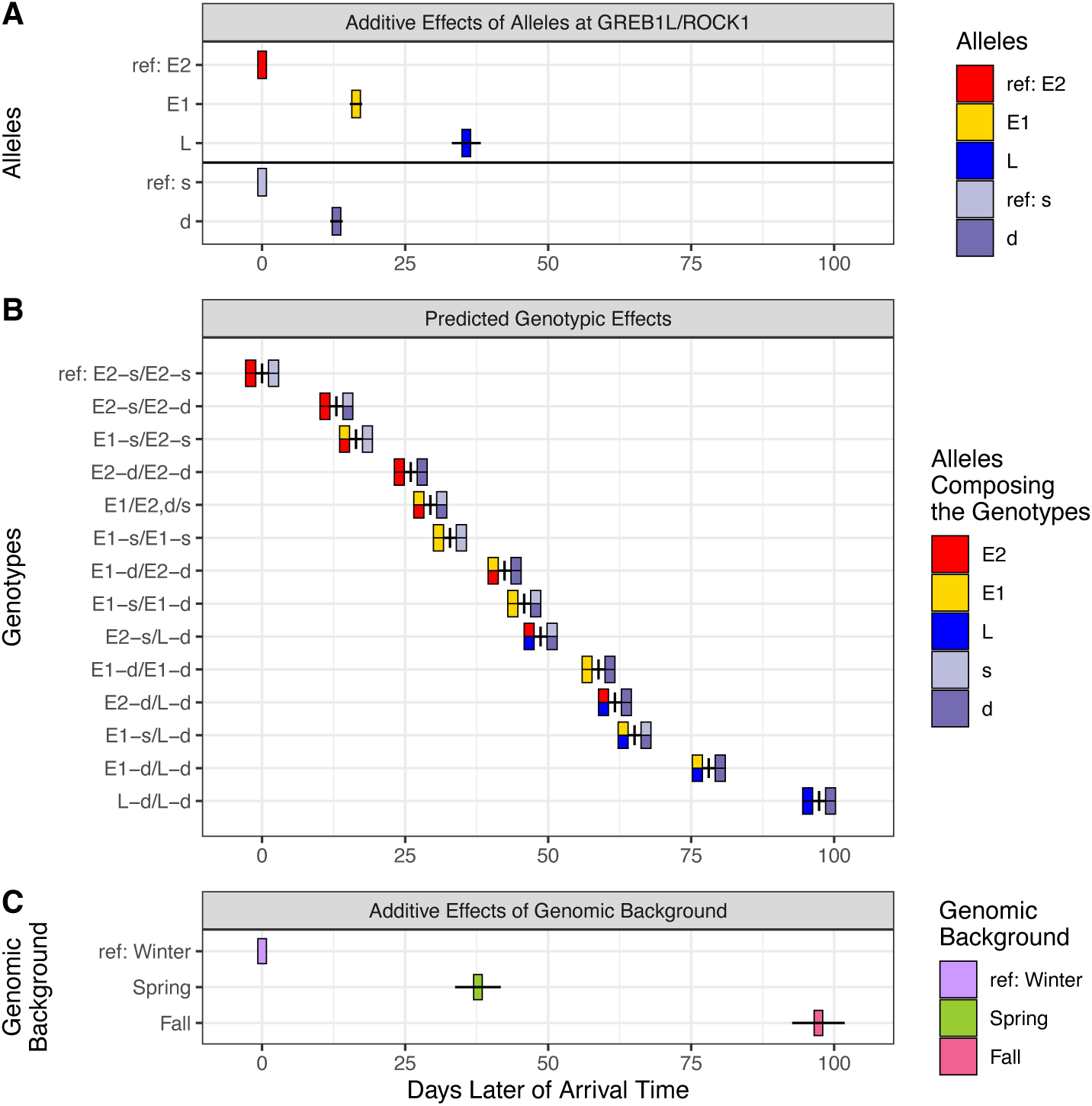
Estimated effects of alleles, genotypes and genomic background. Visual representation of the estimated additive effects of alleles at two domains within the GREB1L/ROCK1 region, their predicted genotypic effects, and the estimated additive effect of genomic background (GB). (**A**) Additive effects of alleles, on the *x*-axis, for different alleles appearing at different heights on the *y*-axis. Each allele is represented by a box centered horizontally on the estimated additive effect. Horizontal black lines atop the center of each box have endpoints at the estimate +/− 1 standard error. In the first domain, (the *E1E2L* domain) the *E2* allele was defined as the reference (effect = 0), while in the second domain (*sd* domain) the non-duplication (*s*) allele is defined as the reference. (**B**) Predicted genotypic effects obtained by summing the additive allelic effects. Different genotypes appear at different heights, are colored according to alleles, and are centered in the *x* direction on their predicted values. (**C**) Estimated effects of GB.

Thus, we show how a single locus of major effect on an important trait can be resolved down at a much finer scale through characterization of functional domains within the locus and with genotypes from large numbers of individuals of known phenotype. The estimated additive effects of the different alleles in these two domains on the predicted adult migration times span over 100 days in fine gradations, providing a remarkable demonstration of the fine-scale variation possible in the critical phenotypic trait for salmon of adult migration timing resulting from only five alleles (Fig. 5B). This provides a vivid example of how continuous trait variation that, in the absence of genetic data, might appear complex and polygenic, can be explained by a relatively simple additive genetic system.

We estimated the effects of the RoSA alleles in a separate sample of 5,508 adult Chinook salmon sampled as they arrived at the FRH in early 2019 (Fig. S5). Genetic assignment revealed that all but two of these fish carry the fall/late-fall GB. The other two fish carried the winter GB and were removed from all further analyses (Tab. S1). The diversity of RoSA genotypes in this sample was considerably lower than in the winter-run fish from KFT (Fig. S5). Nonetheless, applying a truncated regression—due to the evident truncation at the end of the sampling window (Fig. S6)—revealed highly significant additive effects (Tab. S5) for the *E1* and *d* alleles, the two with sufficient observations for accurate estimation. The effect sizes (*E1*: 12.30; *d*: 11.09) were commensurate with those estimated within winter-run fish at KFT.

In the KFT sample, two genotypes, *E1-s/E1-s* and *E1/E2,d/s* (Fig. 3), were found in at least 25 individuals among all three GBs (Fig. 4B). Focusing on these two genotypes we are able to control for the effect of RoSA genotype, allowing us to estimate the contribution of the GB to differences in migration timing. Because there was significant truncation in the sampling of the distribution of migration timing in the fall/late-fall GB (Fig. S7), we again applied a truncated regression and estimated highly significant effects of GB on adult migration timing. The spring GB confers an adult migration time 37 days later, on average, than the winter GB, and the effect of the fall/late-fall GB is 97 days later than the winter GB (Tab. S6). Comparing these GB effects to the predicted effects of different RoSA genotypes (Fig. 5B,C) reveals that the effect of GB is similar in magnitude to that of the RoSA. This provides the first assessment of the relative role of large-effect loci versus the likely polygenic effects of GB on a complex behavioral trait in a free-ranging animal. In contrast, the element of trait variation that is explained by this lineage-specific GB, which is presumably highly polygenic, is not highly variable within lineages and therefore contributes minimally to trait variation within families. That a large fraction of trait variation is apparently due to GB, despite no genomic regions fixed between different GBs, suggests that the migration timing component is likely due to many genes of small effect for which we do not have sufficient data to elucidate contributions (*21*).

Disentangling the effect of GB from that of RoSA genotype is possible because several early-associated genotypes (*E1-s/E1-s* and *E1/E2,d/s*) are present in all three GBs. It is likely that the different RoSA genotypes were historically more strongly associated with different GBs, but have recently been affected by an increase in cross-breeding and associated recombination. For example, it appears that early-associated genotypes have been introduced on the fall/late-fall GB in the Feather River, likely due to extensive crossbreeding of fish of different GBs at the FRH. To confirm that the FRH could have led to such mixing of RoSA genotypes and GBs, we genotyped a representative sample of fish from the FRH fall-run spawning program at our RoSA markers. The results indicate that fish with early-migration-associated genotypes are very often spawned with those carrying genotypes (and hence, historically, also the GB) associated with late adult migration (Fig. S8, Tab. S7, S8). The resulting patterns of genotypic combinations are consistent with extensive crossbreeding between historical late-migrating and early-migrating (both spring-run and winter-run) fish at the FRH, which led to a spring run possessing a fall GB.

We examined the observed date of reproduction (spawning) of fish in the FRH fall-run program with different RoSA genotypes and found that the three genotype categories with sufficient sample sizes to accurately estimate mean spawn dates were the *L-d/L-d*, *E1-s/L-d* and *E2-d/L-d* categories. We found a difference in spawn date of 7.7 days between the homozygote (*L-d/L-d*) and heterozygote categories (*E1-s/L-d* and *E2-d/L-d*), which contrasts with a difference in adult migration timing of approximately 30 days. As with the migration timing trait, there was little difference in spawn timing between the two heterozygote categories, as predicted by the compensatory effects of the two allelic lineages (*E1/E2* and *d/s*). The smaller difference in spawn timing than migration timing for fish homozygous and heterozygous for *E* and *L* alleles is concordant with the pattern observed previously in the Klamath River basin (*8*). This difference has been proposed to be a consequence of the acceleration of reproductive maturation due to the increased time spent in warmer freshwater by early-migrating fish (*8*).

We note that there is a contrast, however, in the CCV and Klamath River basin with respect to the association of RoSA genotypes and GB. Whereas the Klamath River populations have genomic relationships based entirely on geographic distance (*22*), in the CCV, there are distinct GBs associated with populations that occur parapatrically in the same subbasins but were historically dominated by either late-(*L*) or early-migrating (*E1*, *E2*) RoSA genotypes (*15*). For example, the spring-run fish from Butte, Mill and Deer creeks still carry almost exclusively *E1* alleles (Fig. 4A) and have a GB distinct from fall-run fish in those subbasins, whereas in the Klamath individuals from a local population share the same GB regardless of their RoSA genotype.

A previously identified genomic region with large allele frequency differences between late-fall-run fish and all other CCV Chinook salmon, termed the Late-Fall-Associated Region (LFAR, (*15*)), did not yield a statistically supported association with migration timing (Fig. S9). However, we note that the lack of individuals homozygous for non-LFAR alleles and with late-fall-run migration timing in our dataset means we can not confidently rule out a dominant effect of this region on migration timing. Moreover, our sampling strategy does not include trapping of fish during the seasonal period (Oct.–Dec.) when the more abundant fall-run fish migrate, and lack of sampling during these months limits our ability to detect an effect of this locus on migration timing. However, we note that the proportions of the sampled genotypes are not different from Hardy-Weinberg expectations (chi-square test, *p* = 0.28, 0.32, for the two SNPs), indicating that the sampling period is likely representative of the ecotype. Regardless, the lack of association at this locus with migration timing indicates that differences of the late-fall-run ecotype are unlikely to be due to a single locus of major effect and more likely due to polygenic factors. Such factors may be due to selection acting on elements of ecotypic life history other than adult migration timing.

Similarly additional genomic variants beyond the GRR with large allele frequency differences between winter-run fish and all other salmon in the CCV have been previously described and termed the Winter-Run Associated Polymorphisms [WRAP; (*15*)]. We investigated the WRAP markers in our samples and found no statistically supported association with adult migration timing within different ecotypes. We note that this indicates, as with the LFAR, that the ecotypic differences of winter-run associated with the GB are likely polygenic. Moreover, the observed allele frequency differences may be a consequence of selection on other elements of the winter-run life history or physiology, including different migratory pathways in the ocean or different thermal tolerance thresholds.

We found a striking pattern of genotypic distribution, with only three copies of the late (*L*) haplotype present (i.e., a frequency of 0.0003), all in heterozygous form, in the near-complete sampling of the spring-migrating salmon in the Feather River in 2019. Earlier work in the Klamath River found that RoSA heterozygotes primarily migrated and spawned during the spring-run (early-migrating) distributions, even though they had an intermediate mean timing of adult freshwater entry (*8*). The lack of heterozygotes in the spring-run arrivals in the Feather River of the CCV indicates different dominance patterns of the RoSA in the CCV and the Klamath River basins. We also note that heterozygotes sampled in the CCV in that previous work were all field-identified as fall-run salmon, due to the timing of spawning (*8*). We note, however, that this lack of heterozygotes in the spring-arriving fish is not due to their absence in this population. They are numerous, but apparently have later migration timing than *EE* homozygotes, and spawn during the fall-run reproductive period, but with a slightly earlier mean spawn timing (Fig. S8, Tab. S7, S8). This is in contrast with the Klamath River, where heterozygotes all spawn in the early-migrating (spring-run) reproductive period (*8*).

Thus, the available evidence indicates that *EL* heterozygotes in the CCV have a predominantly late-migrating life history. We note however, that our study design is not capable of estimating dominance effects of the *L* allele (because of the lack of sampling of *LL* homozygotes at KFT and of summer sampling in the Feather River). The mechanism responsible for this apparent shift in the dominance relationship is not clear, but could be due to a distinct modifier locus found in the CCV, or some other element of the GB. Alternatively, here we measure the freshwater migration of fish to the terminus of anadromous access, and not the exact timing of entry into freshwater, so it is possible that this is a reflection of the different sampling strategies in these studies. However, in the Klamath River, heterozygotes mostly spawn in the early-migrating (spring-run) reproductive period.

Previous work (*6*, *7*) found that variants in the RoSA were associated with adult migration timing in two species of related salmonids, leading some authors to suggest that this region “controlled” the phenotype. They further suggested that this implied a shared mechanism in the two species which was most likely acting through the reproductive maturation pathways. However, subsequent work showed that the seasonal shift in life history is entirely in the migration pathway, at least in Chinook salmon, with reproductive maturation following a similar schedule regardless of migration timing (*8*). That work also suggested that the RoSA was a simple two-allele system that had largely non-overlapping trait distributions for the two homozygous genotypes. While that may be the case in the original study system, the Klamath River basin in Northern California/Southern Oregon, we show here that adult migration timing of Chinook salmon in the CCV is more complex and that the RoSA acts as a five-allele system that generates a highly gradated distribution of migration timing. Similarly, Horn et al. (*11*) found a functional effect of the intergenic duplications on migration phenotype in Columbia River Chinook salmon, indicating that such complexity in the effect of the RoSA on salmon life history may be widespread.

Closer inspection of the intergenic duplications (*sd*) region of the genome at http://genome.ucsc.edu (*23*) revealed that this region spans, and is bordered by, several CpG islands. CpG islands often appear near the beginning of transcription regions in some vertebrates and may be associated with promoter regions (*24*). It is possible that the increase in length of this intergenic region, due to the duplications, may impact expression of genes that affect this complex trait. More research is required to elucidate the functional elements of this genomic region.

The effect of GB on migration timing in the CCV appears to differ from that in the Klamath River basin. In the Klamath basin, significant population structure exists, with differentiation between the most distant geographic locations similar in magnitude to the values between the ecotypically defined Evolutionarily Significant Units (ESUs) in the CCV (*22*). However, in the Klamath River, structure follows the expectations of an isolation by distance model, with a strong correlation between geographic and genetic distance, which indicates that migration between proximate populations is the primary determinant of differentiation. Conversely, in the CCV, the ecotypes and associated GBs appear to have evolved in allopatry or in parapatry. As such, early-and late-migrating fish from each of the two main populations in the Klamath River basin share a GB, regardless of adult migration phenotype. Moreover, even though there are clinal differences in GB within the Klamath River basin, they do not appear to affect the seasonal migration timing phenotype, and the GRR is the sole determinant of seasonal migration timing in that basin (*8*).

Previous work in the Klamath River found that the *E* haplotype was partially dominant, with the migration and spawn timing of heterozygotes overlapping more with the early-migrating (spring-run) fish than the late migrating ones. Moreover, in locations where access to cold-water habitat in summer had been eliminated (*8*, *25*), as in the upper Klamath basin, this partial dominance facilitated the loss of the *E* haplotype as heterozygotes were subject to similar selection against summer residence.

In contrast, in the CCV, it appears that the L haplotype is partially dominant, with very few heterozygotes migrating early, but many heterozygotes observed in the late-migrating spawners (Fig. S8). While we lack exact migration timing information for CCV RoSA heterozygotes, it is clear that the *E* haplotype is more likely to be maintained in CCV populations during periods of limited cold-water habitat in summer, perhaps providing additional resilience for the life-history variation in this population complex. Given the highly variable nature of climate and stream phenology in California, this will provide a modest buffer that may facilitate future recovery of robust early-migrating salmon populations.

The winter-run ecotype descended from a common ancestor with the CCV spring-run lineage (*17*). As such, and given that it is absent elsewhere in the species range, the E2 lineage appears to have evolved in the winter-run group. It thus represents a globally distinct allele that contributes quantitatively to the local adaptation that is necessary to access the highest-elevation Chinook salmon habitat in the region, and is one of the first instances of a molecular-level description of a unique adaptation of a rare and endangered species.

We show that a simple two-locus, five-allele system in the GRR, in conjunction with the evolutionarily distinct GBs associated with the two early-migrating lineages in the basin (winter run and spring run), produces an almost continuous stream of fish entering the CCV. We provide a mechanistic explanation for the well-known distinction of the Sacramento River basin as being the only basin range-wide with adult fish present year-round (*26*). Understanding the genomic architecture of migration timing in Chinook salmon is a critical first step in constructing a predictive model for how changes in anthropogenic activities, such as hatchery production and river flow management, will affect the distribution of trait values for this key behavior. That, in turn, will provide a framework for designing conservation and management strategies that will meet societal and legal goals for the maintenance and recovery of depressed salmon populations. For example, it may be feasible to use information about allelic variation in a marker-assisted selection program to maintain particular phenotypic optima in populations of salmon that are supplemented with hatchery-produced fish. This would allow better phenotype matching with predicted environmental conditions and the structure of fishery seasons. It may also allow the maintenance or even resurrection of distinct populations of salmon that are able to best utilize habitat that becomes newly available through actions such as dam removal, or in the face of climate change-altered phenology. Moreover, the association of variation in migration-timing phenotypes with that of other life-history traits, such as ocean migration routes and age at maturity, indicates that such a predictive framework will also serve to maintain more complex salmon biodiversity in the CCV and further afield.

## Material and Methods

### Genetic Markers, reference populations, and population differentiation

For genetic stock identification (*27*) and population assignment (*28*) of migratory fish, we genotyped a reference or “baseline” sample of 1,073 Chinook salmon encompassing nine different populations from seven CCV locations (Fig. 1) encompassing winter-, spring-, fall- and late-fall-run fish. We used a previously published (*8*) set of 125 microhaplotype (*29*) loci (Data S1) and assessed the accuracy of this reference data set for population assignment via leave-one-out cross validation using the R package ‘rubias’ (*30*). A small number, 32, of fish likely belonging to populations from which they were not sampled (being either migrants or sampling errors) were evident in the genetic data, most conspicuously in fish sampled from Butte Creek spring (BCS) and Feather River Hatchery spring (FRHS) that were assigned to the winter genomic background (GB). Since the winter run is so distinct from all other populations, and they are known to occasionally intermingle with other early-migrating fish (*15*), these “misassignments” are clearly just winter-run fish incorrectly sampled within BCS and FRHS collections. The same is likely true for several of the fish in Mill/Deer Creek spring (MDS) that assign to fall-run populations (i.e., some are fall-run fish that happened to be sampled and categorized as spring run in MDS). Nonetheless, to be conservative, we retained all fish in the accuracy measurements. The same genotype data set was then used to evaluate *F*_ST_, a measure of genetic differentiation, between the nine populations. Additionally, six of those nine populations were included in a previous study (*8*), and the whole-genome sequencing data from that study was used to estimate pairwise *F*_ST_ between those six populations using ANGSD (*31*) in combination with winsfs (*32*).

To supplement the microhaplotype markers with amplicons that identify the *L*, *E1*, and *E2*, as well as the *s* and *d* alleles, we started from the eight SNPs published in (*8*), and then identified four more to add to those (Tab. S2, Data S2). The four new SNPs were found by first surveying 13 SNPs with alleles that were fixed, or nearly fixed, on the *E1* and *E2* haplotypic backgrounds (Fig. S1) in the whole genome sequencing data of (*8*). Of these 13 SNPs, three SNPs were chosen to identify the *E1* and *E2* haplotypes. Additionally, we added a single SNP which we call RA7.T, previously reported as snp670329 (*25*), to identify *s* and *d* alleles carried in the region of the intergenic duplications. In order to establish that the two alleles at this locus are highly associated with the presence or absence of the duplicated segments in the region between the *GREB1L* and the *ROCK1* genes, we expanded upon the work in (*8*) which first defined the genomic ranges that contained segments that were potentially duplicated, and then compiled the observed sequencing read-depths within and outside of those potentially duplicated segments within the intergenic region, for each individual, who were themselves categorized according to whether they were homozygous or heterozygous for an *L*-lineage or *E*-lineage haplotype. We continued that analysis by comparing the bases observed at the genomic position of RA7.T in all fish sequenced in (*8*) that had at least one read covering RA7.T. We used the samtools (*33*) mpileup subcommand to extract these bases. By comparing the observed bases (different colored points) with the read depth observed versus that expected throughout the potentially duplicated segments amongst spring run (few of whom carry the duplications), the correspondence between allele at RA7.T and the presence of duplications can be made clear.

### Samples of adult, migrating salmon

We obtained samples of migrating, adult Chinook salmon from two locations in the California Central Valley (CCV): the Keswick Dam fish-trapping facility and the Feather River Hatchery (Fig. 1).

Keswick Dam is a 49 m high dam on the mainstem of the Sacramento River at the terminus of anadromy for the basin, 15 km downstream from Shasta Dam and 486 km upstream from the Sacramento-San Joaquin Delta. It houses a fish-trapping facility that typically operates from the last week of December, through the winter, spring, and summer until August 1. This period coincides with the arrival times at Keswick Dam of three distinct ecotypes, or “runs,” of Chinook salmon: the late-fall run (late December through late March), winter run (late January through July), and spring run (early April through August). Trapped fish are either transported to one of several hatcheries for spawning, or are released downstream to spawn in tributaries below Keswick Dam.

Between the beginning of August and the end of December, the Keswick Fish Trap (KFT) does not operate consistently. This period coincides with the migration time of the fall-run Chinook salmon, the most abundant ecotype in the CCV. We received tissue samples and their arrival/sampling dates, from 5,741 Chinook salmon handled at the KFT during the years of 2009 to 2018.

The Feather River Hatchery was established in 1967 to mitigate habitat made inaccessible due to the construction of Oroville Dam on the Feather River. It is located at the base of Oroville Dam, at the terminus of anadromy for the Feather River, one of the largest tributaries of the Sacramento River (Fig. 1). The hatchery propagates both spring-and fall-run Chinook salmon. Starting in 2003, a broodstock selection protocol was established to minimize cross-spawning of fall-run fish in the spring-run breeding program at the hatchery: any fish ascending the ladder to the hatchery before July 1 of each year is marked with an externally visible tag to indicate a spring-run fish and released back to the river. In 2019, the arrival date of each of these tagged fish was recorded, and tissue for DNA extraction was collected from 5,631 fish before they were released back to the river. For almost three months after July 1, fish are not allowed to enter the hatchery, and they remain unmarked in the river. When entry to the hatchery is reopened in late September to collect fish for spawning as broodstock in the spring-run program, externally tagged fish are mated only with other such fish (the putative spring-run spawners). After about one month, many fewer externally tagged fish migrate into the hatchery. As such, in mid-October the FRH ceases to use externally tagged fish as broodstock and uses only fish that did not enter the hatchery in spring, as spawning broodstock for the fall-run program.

There is spawning and holding habitat downstream of the dam/hatchery and not all salmon that migrate into the Feather River enter the hatchery during either the early (spring) or late (fall) migration periods. As such, not all spring-run salmon that migrate into the Feather River in the early migration period will receive external tags.

To investigate the distribution of *L*, *E1*, *E2*, *s*, and *d*, alleles amongst the fish used in the fall-run spawning program at FRH, we obtained tissues from the 3,284 fish spawned over 16 days in the fall-run program in spawn year 2008. From these, we genotyped 768 fish, with roughly 50 fish (when at least that many were available) randomly sampled from each day of spawning.

The sampling schedules at each hatchery facility are tailored for specific management purposes, which affects the inferences that are possible. Most importantly, there is the possibility for truncation of data. Because the KFT does not typically operate after August 1, fish arriving after that date are not sampled; while most of the unsampled fish are believed to be from the fall run, some spring-run fish may arrive after August 1 and are not sampled at the KFT. This type of data truncation is also possible at FRH. The analyses we choose for each data set reflect this.

### Analysis of haplotypic variation in the RoSA region on migration timing

We first investigated patterns of linkage disequilibrium between the 12 markers of our RoSA marker panel amongst the KFT and FRH migrant samples to establish that our panel of 12 SNPs readily identifies haplotypic structure in the region. To do this, we used a simple approach (*34*), whereby we first identified haplotypes that are known without uncertainty due to the fact that they occur in individuals that are homozygous for all 12 markers. Subsequently, when possible, we resolved heterozygous individuals into pairs of haplotypes that were previously identified in homozygous form. This approach identified two separate domains spanned by our markers, one with three haplotypes/alleles, *L*, *E1*, and *E2*, (hereafter the *LE1E2* domain) amongst our samples, and the other, the *sd* domain, with two alleles, *d* and *s*, corresponding to chromosomes that do contain and do not contain the duplication, respectively (See Results and Discussion).

For fish sampled at the KFT, we used the 125-locus reference baseline of populations to assign individuals to one of the three reporting units: Winter, Spring, and Fall/Late-Fall. Because the KFT sampling does not encompass the migration time of the fall-run ecotype, individuals assigned to this last reporting group are presumed to be either late-fall-run or Feather River-lineage spring-run fish, depending on their RoSA genotypes. These reporting units correspond to the three different GBs of fish that can be readily distinguished via population assignment with the 125-locus reference baseline. After assigning the KFT samples to these three lineages, we visually explored, by plotting points and boxplots using the ‘ggplot’ package (*35*) in R, the relationship between timing of migration (measured as date of sampling at the KFT), and lineage, as well as genotype at the *LE1E2* and *sd* domains. From these visualizations we saw that a small number of salmon had been sampled outside of the standard sampling period of late December to August 1; such fish were removed from further analysis.

From the data visualization, it was evident that the winter-run lineage had the most diversity in *LE1E2* and *sd* genotypes (see Results), making it a suitable candidate for estimating the effect upon migration timing of different alleles in those two domains. We modeled the effects of the *LE1E2* and *sd* domains as additive in a linear model. Within each subdomain we chose a natural and orthogonal interactions, or NOIA, approach for estimating additive and dominance effects of the different alleles (*36*). For the tri-allelic *LE1E2* domain, following (*37*), we designated the *E2/E2* genotype (which is the earliest returning genotype at the *LE1E2* subdomain) as the reference, and constructed the design matrix with all additive and dominance effects as follows:

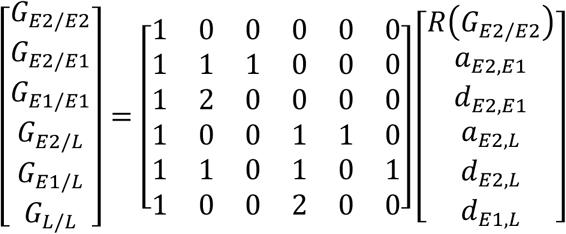

The left-hand side of the equation holds the vector of genotypic effects on migration timing with *G_x/y_* indicating the overall effect of a genotype consisting of alleles *x* and *y*. The column vector on the right side holds the parameters to be estimated, as follows: *RG_E_*_2/*E*2_/ denotes that the genetic effect of the *E*2/*E*2 genotype is being used as the reference, *a_x,y_* denotes the additive effect from substituting a *y* allele in place of *x*, and *d_x,y_* is the dominance effect of a *y* allele when in heterozygous form with an *x* allele. For the biallelic *sd* domain, we designated the *s* allele (the allele associated with earlier adult migration) as the reference in the NOIA design:

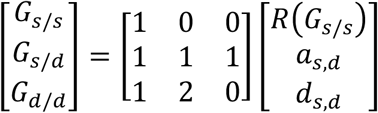

Within the winter-run lineage, we included these genetic models for the two subdomains as fixed effects with year as an additional covariate in a series of linear models, defining migration date at Keswick *D_K_* as the number of days each year since the start of sampling on Dec 26 that each fish was encountered and sampled. We first considered year as a random effect in a linear mixed model:

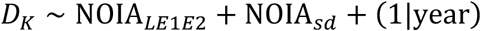

We also considered a similar model with year as a fixed effect,

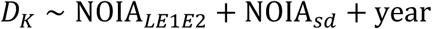

which we fit by linear regression, and also using a truncated regression model, using the ‘truncreg’ package (*38*) in R, to see if there was evidence that some fraction of the later-arriving winter-run *GREB1L*-region genotypes remained unsampled, since they arrived after August 1.

Finally, the linear-mixed model analysis was repeated for the winter run at KFT with the relatedness between individuals accounted for by including the genomic relationship matrix (GRM) calculated from the 125-locus microhaplotype data set as a random effect using the R package ‘sommer’ (*39*). Since our data is multiallelic, we calculated the GRM using the approach taken by (*40*), as adapted by (*41*) for using multiallelic haplotypes. Briefly, each microhaplotype allele at a locus is treated as if it were an allele at a biallelic locus, and individuals are coded as a 0, 1, or a 2, according to how many copies of the allele are present in the individual. More specifically, let *I_i_*_ℓ*k*_ count the number of alleles of type *k* at locus ℓ within individual *i*, and let *p*_*ℓk*_ denote the relative frequency of allele *k* at locus ℓ in the sample from the population. Missing values of *I*_.ℓ*k*_ are replaced with their expectation, 2*p*_ℓ*k*_. Then, define the centered variables, *X_i_*_ℓ*k*_ = *I_i_*_ℓ*k*_ − 2*p*_ℓ*k*_. If we let *j* index the unique combinations of ℓ*k*, then we can write that as *X_ij_* = *I_ij_* − 2*p_j_*, and these can be organized into an *N* × *K* matrix **X**, where *N* is the number of individuals (i.e., *i* = 1, …, *N*) and *K* is the total number of different alleles across all loci (*j* = 1, …, *K*). The GRM, **G**, is then

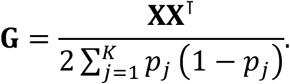

### Analysis of RoSA-region effects on migration timing at FRH

The fish returning to FRH, whether they are part of the spring-run or the fall-run program, are all of one GB, the fall/late-fall. There were only four genotypes at the *LE1E2* and *sd* domains, together, with more than seven observations: *E1-s/E1-s*, *E1/E2,d/s*, *E1-s/E1-d*, and *E2-d/E2-d*. These were analyzed using the NOIA framework with reference genotypes being *E2/E2*, and *s/s*. Because there were no *L* alleles found in the FRH sample, and because the dominance effect at the *sd* domain was not estimable with only the four genotypes occurring in appreciable numbers, we used natural restrictions of the above NOIA models for the *LE1E2* domain (NOIA^↓^ *_E_*_1*E*2_):

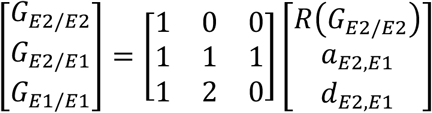

and for the *sd* domain (NOIA^↓^ *_E_*_1*E*2_)

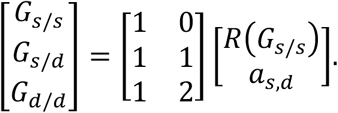

With *D_L_* being the date that the fish was encountered and sampled at the FRH fish ladder, the model is, then,

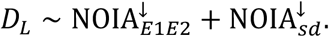

We used a truncated regression implemented in the ‘truncreg’ R package (*38*) to account for the considerable truncation evident in the histograms of arrival time (Fig. S7) at the FRH ladder.

### Genomic lineage effects on migration timing at KFT

Only two genotypes (*E1-s/E1-s*, and *E1/E2,d/s*) were found in sufficient numbers in all three genomic backgrounds (winter, spring, and fall/late-fall) for an analysis of the effect of GB. We thus restricted the data set to these two genotypes, and we modeled each genotype as a discrete category. Our first model was a linear mixed model with year, grouped by lineage, as a random effect:

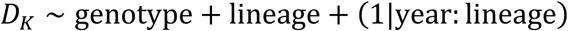

Subsequently we investigated the linear model with year as fixed effect,

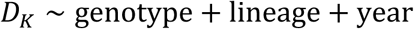

and analyzed that, as well, using truncated regression.

### Effect of LFAR and WRAP

To investigate potential associations between the LFAR (Late-Fall Associated Region) and the WRAP (Winter-Run Associated Polymorphisms) markers and adult migration timing in Chinook salmon, we examined genotype data from individuals collected at the KFT. We genotyped all fish from the KFT with two SNP markers at locations 954,054 and 1,151,868 on Chr34 in the Otsh_v2.0 version of the Chinook salmon genome (GenBank Accession: GCA_018296145.1) that were previously described as having large allele frequency differences between late-fall run and all other fish in the CVV (*15*). Similarly, we genotyped all fish from KFT at three previously described markers (at locations 27,450,181 on Chr08, 70,794,116 on Chr12 and 36,409,831on Chr16 in the Otsh_v2.0 genome with large allele frequency differences between the winter run and all other fish in the CCV (*15*) For the LFAR markers, we analyzed late-fall individuals (N = 88), categorized as described above, and for the WRAP markers, winter-run individuals (N = 1,203). In each case we modeled adult migration timing with a linear mixed model identical to the model for the *LE1E2* and *sd* domains, though expanded with a fixed NOIA term for each of the markers being tested.

### GREB1L-region genotypes amongst the FRH fall-run-program spawners

The 768 samples were genotyped at the 12 markers in the *LE1E2* and *sd* domains. Keeping only the individuals that had genotype calls at all 12 markers, we summarized the data by designing a genotype-bar plot that shows the number of fish with different genotypes at the two domains on each day of spawning. There was variation each day in the fraction of spawners that were subsampled and successfully genotyped. If *N*_t_ fish were spawned and *n*_*t*_ were genotyped on day *t*, that fraction is *f*_*t*_ = *n*_*t*_/*N*_t_. To account for that varying fraction, each day, we also plot the results in a genotype-bar plot in which the height of each individual sample has been expanded by a factor of 1/*f*_*t*_ to indicate the number of spawned fish that it represents.

## Acknowledgments

We thank staff from the US Fish and Wildlife Service, the California Department of Water Resources, and the California Department of Fish and Wildlife for assistance in the collection of samples used in this work, particularly K. Niemala, K. Offill, C. Smith, J. Kindopp, other CDFW/DWR staff at the FRH, and D. Pearse, S. Edmondson and E. Correa at NOAA and UCSC. USDA is an equal opportunity provider, employer, and lender. Mention of trade names or commercial products in this publication is solely for the purpose of providing specific information and does not imply recommendation or endorsement by the U.S. Department of Agriculture.

## Author Contributions

Conceptualization: ECA, NFT, JCG

Methodology: ECA, AJC, NFT, JCG

Investigation: ECA, AJC, NFT, AKB, EC, CC

Visualization: ECA, AJC

Supervision: ECA, JCG

Writing—original draft: ECA, JCG

Writing—review & editing: all authors

## Funding Information

Califorina Sea Grant Fellowship #UCSD 18768 to NJT

** Data Availability**

Raw data and analytical resources used for results will be made available on GitHub/Zenodo, when accepted.

## Authors Declare no Competing Interests

**Fig. S1:**
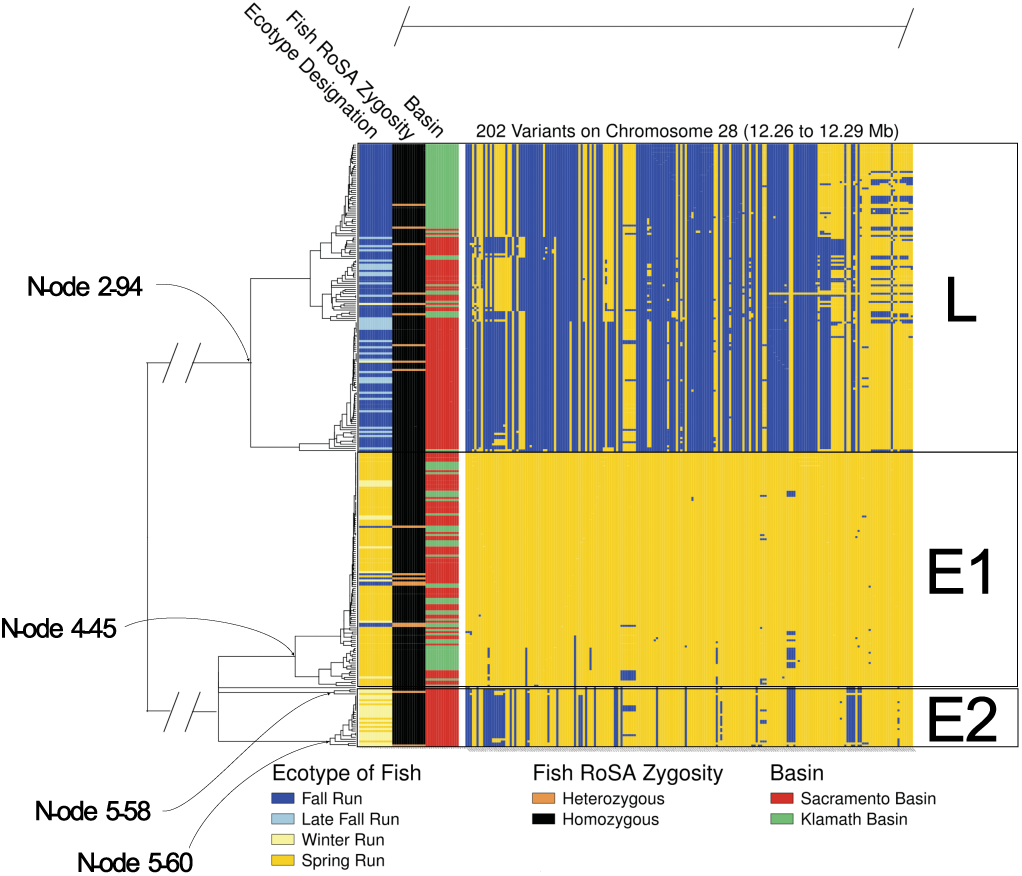
A haplotype raster and neighbor-joining tree explicitly showing the *L*, *E1* and *E2* haplotypes. Within the raster each row is one of two haplotypes from an individual, and each column is a variant. Yellow cells denote an allele most frequent in the spring-run, while blue denotes an allele that is least frequent in the spring run. A version of this figure appears in (*8*). Here, the nodes of the tree subtending the different haplotypes are denoted, and the groups of *E1*, *E2*, and *L* haplotypes are illustrated.

**Fig. S2:**
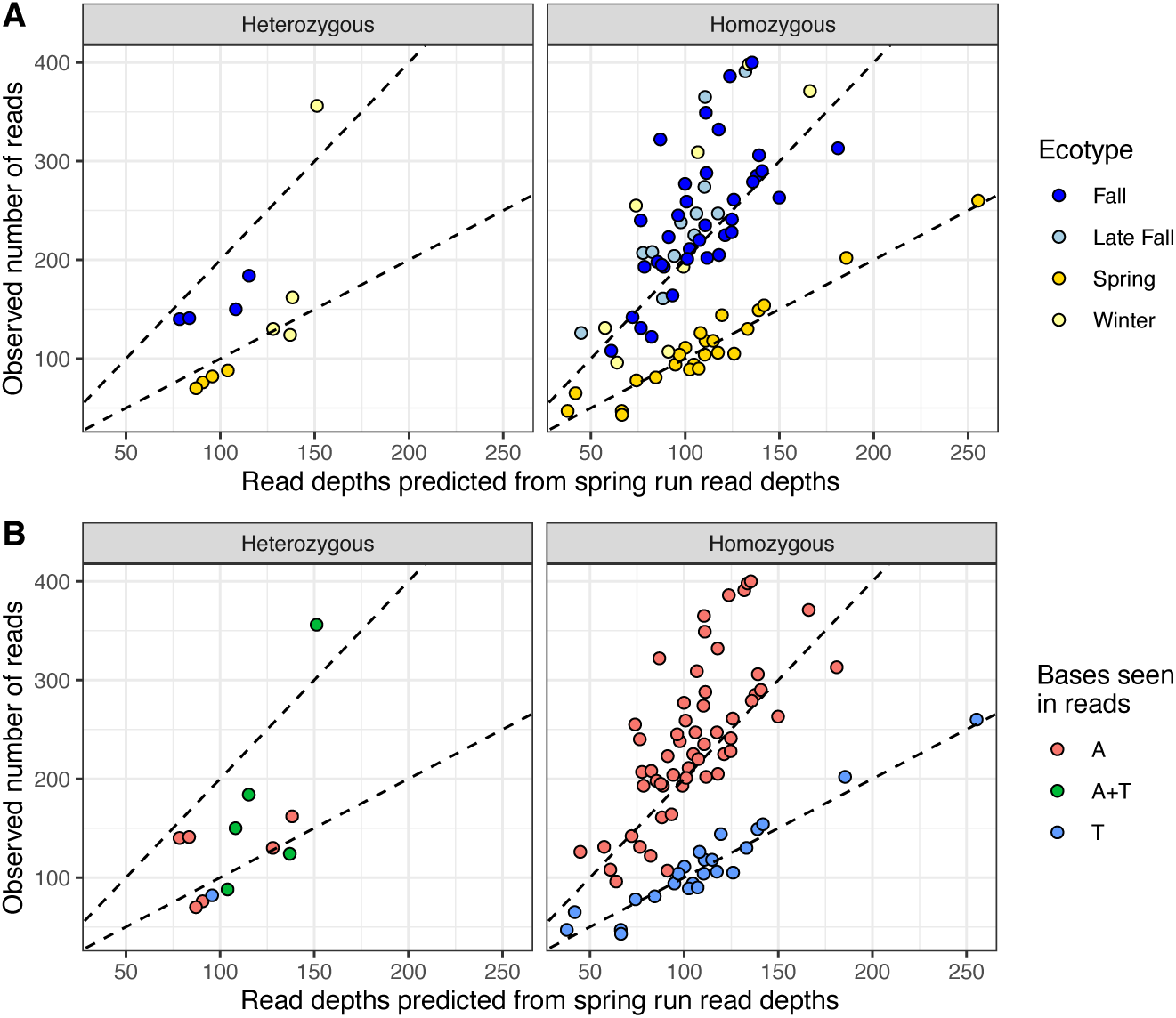
SNP RA7.T tags the presence of the intergenic duplications in the GRR. Panel (**A**) This is a re-rendering of Fig. S6 in (*8*)—the plot of read depths from the genomic regions that are subject to duplication, with points colored according to nominal ecotype. The version here is restricted to fish having at least one read covering the position of RA7.T, and the data have been plotted in two frames corresponding to whether the fish carried two copies of either an *E*-lineage haplotype or an *L*-lineage haplotype (Homozygous) or one copy of each (Heterozygous). The *x*-axis is the number of reads from the region subject to duplication, predicted for spring-run fish from each fish’s average, genome-wide read depth. The *y*-axis is the observed number of reads in the region subject to duplications. Shown on the plot are the *y* = *x* line, around which fish not carrying the duplications cluster, and the *y* = 2*x* line, around which fish carrying the duplication cluster. (**B**) The same fish plotted as in (**A**), but colored according to which bases were observed in sequencing reads at the RA7.T position. The clustering indicates a nearly perfect correspondence, with the T and A nucleotides signifying the absence and presence of the duplications, respectively.

**Fig. S3:**
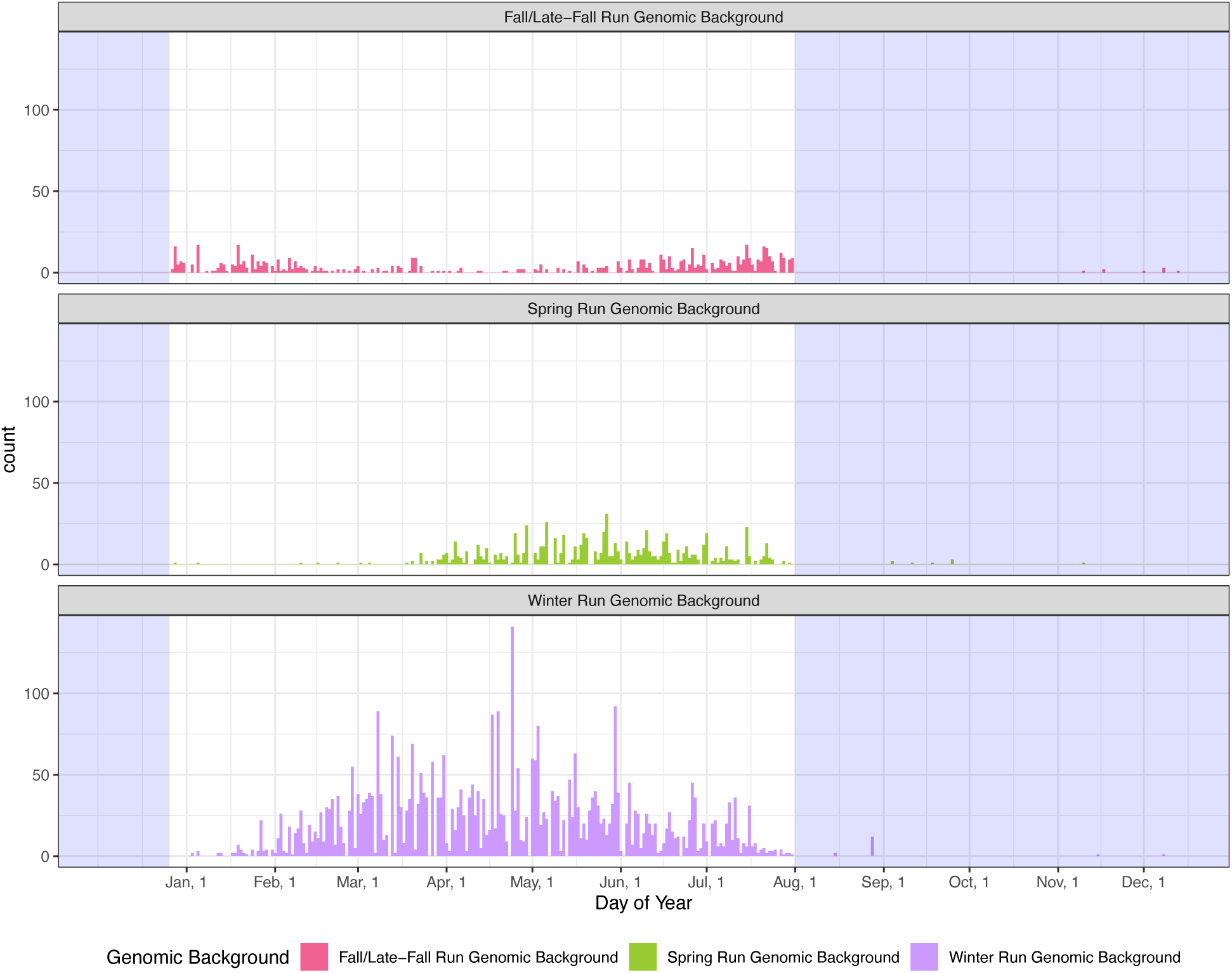
Distribution of run timing across all samples at KFT. The *x*-axis shows adult migration time (sampling date at KFT) across all years of data for all fish in the sample. Fish assigned to the different genomic backgrounds with the 125-marker genetic panel are indicated by different colors and in the different panels. The sampling window at KFT starts December 26 and ends on August 1 of the following year. Times outside the KFT sampling window are indicated by the blue-shaded regions of the plot. Samples outside the sampling period were discarded as many occurred in only one or several years and were likely metadata errors or were samples of fish taken at the hatchery during or before spawning at staff training days and included in the data set. Note that the winter and spring genomic backgrounds appear to suffer little truncation by the August 1 end-of-sampling date, while significant truncation is likely in the fall/late-fall genomic background.

**Fig. S4:**
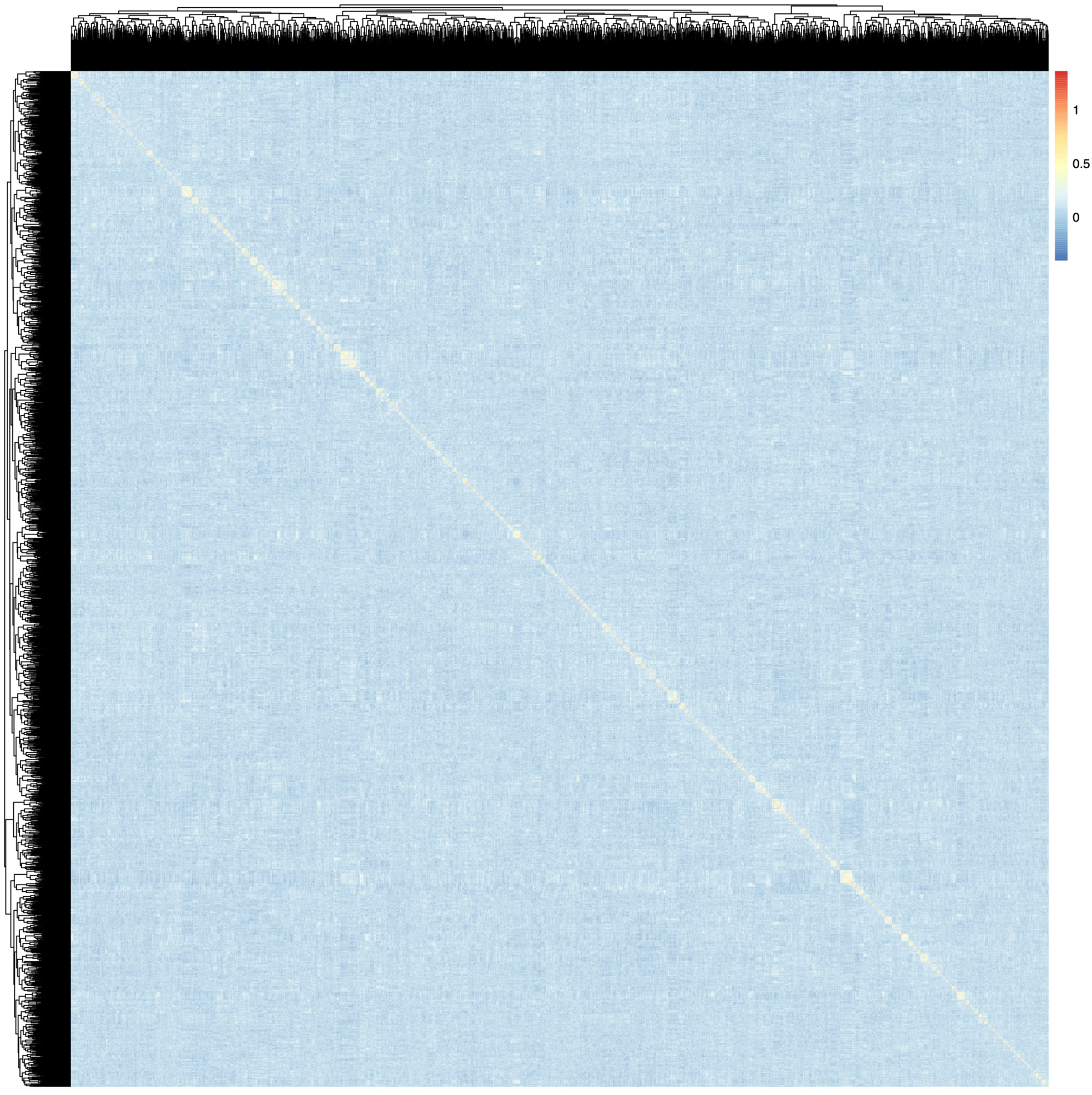
Genomic relationship matrix of winter-run fish at KFT. The GRM was calculated using the 125-locus microhaplotype panel. Fish missing data at more than 28% of the loci were excluded from analysis, leaving 3,000. Each cell in the image represents the relatedness estimate between two individuals. There are 3000 rows and columns that have been hierarchically clustered to place related clusters together on the diagonal. The yellow-hued blocks along the diagonal indicate full-sibling groups (sometimes clustered within half-sibling groups).

**Fig. S5:**
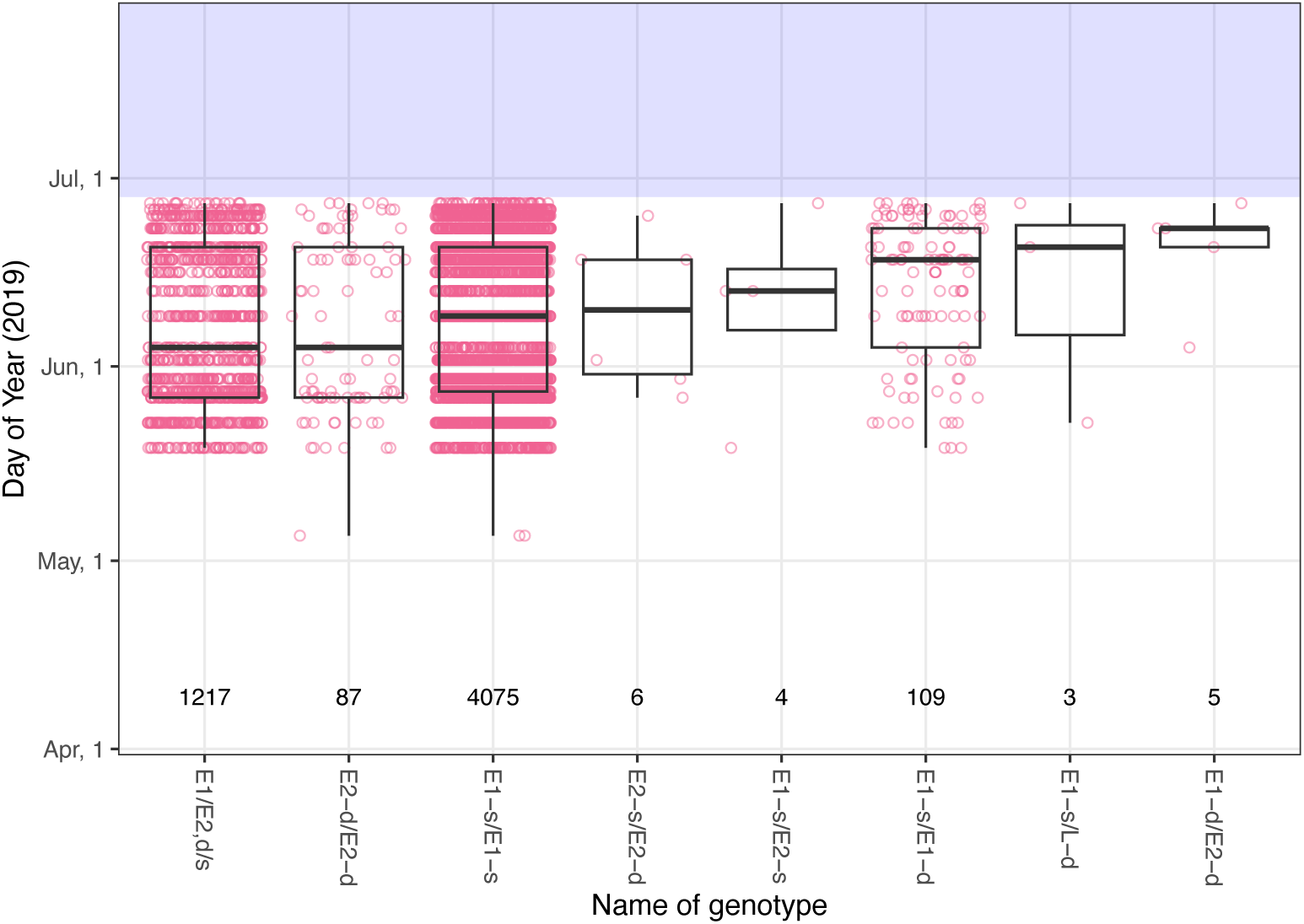
Adult migration timing of FRH spring-run program spawners. The *x*-axis shows different RoSA genotypes, the *y*-axis gives date of arrival and sampling at the FRH fish ladder. All these fish were assigned to the fall/late-fall GB. We include only the four genotypes with more than 6 samples in a truncated regression to determine additive effects of the *E1* and *d* alleles. The transparent blue rectangle represents the end of the sampling period, which suggests some truncation of samples (Fig. S6).

**Fig. S6:**
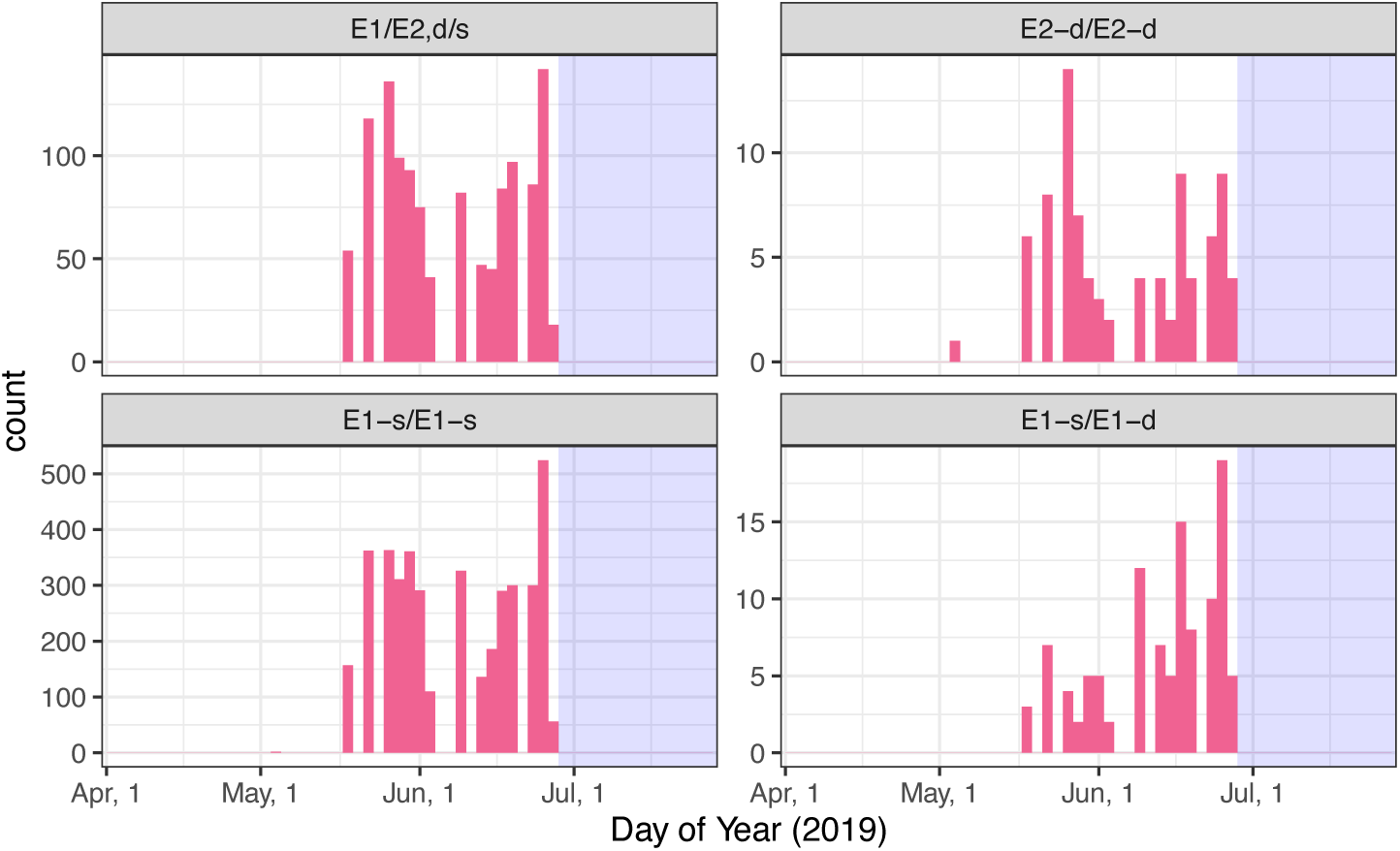
Likely truncation of adult migration timing of FRH spring-run program spawners. Distribution of sampling date at the FRH Ladder for four genotypes included in the analysis of adult migration timing. The sampling window closed June 29, 2019 (blue rectangle), and the distribution of values suggests this may have truncated data, especially in the *E1-s/E1-d* genotype.

**Fig. S7:**
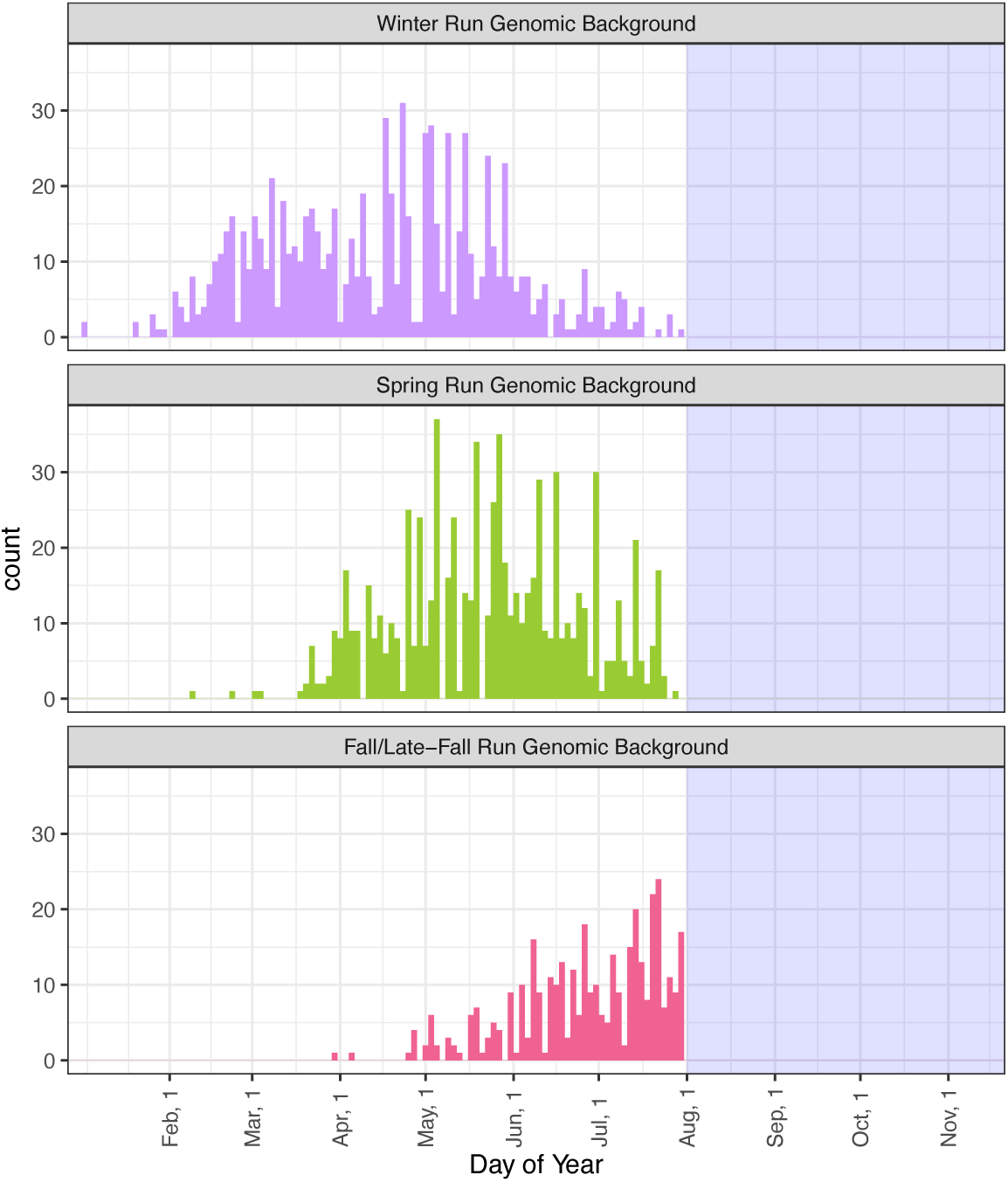
Run timing at KFT of *E1/E2,d/s* and *E1-s/E1-s* genotypes in three genomic backgrounds. The *x*-axis shows adult migration time (sampling date at KFT) across all years of data for all fish carrying the *E1/E2,d/s* or *E1-s/E1-s* genotypes at the GRR. Fish assigned to the different genomic backgrounds with the 125-marker genetic panel appear in the different panels. The sampling window at KFT starts December 26 and ends on August 1 of the following year. Times outside the sampling window are indicated by the blue-shaded regions of the plot. Note the profound truncation evident in fish from the fall/late-fall run genomic background carrying these two genotypes.

**Fig. S8:**
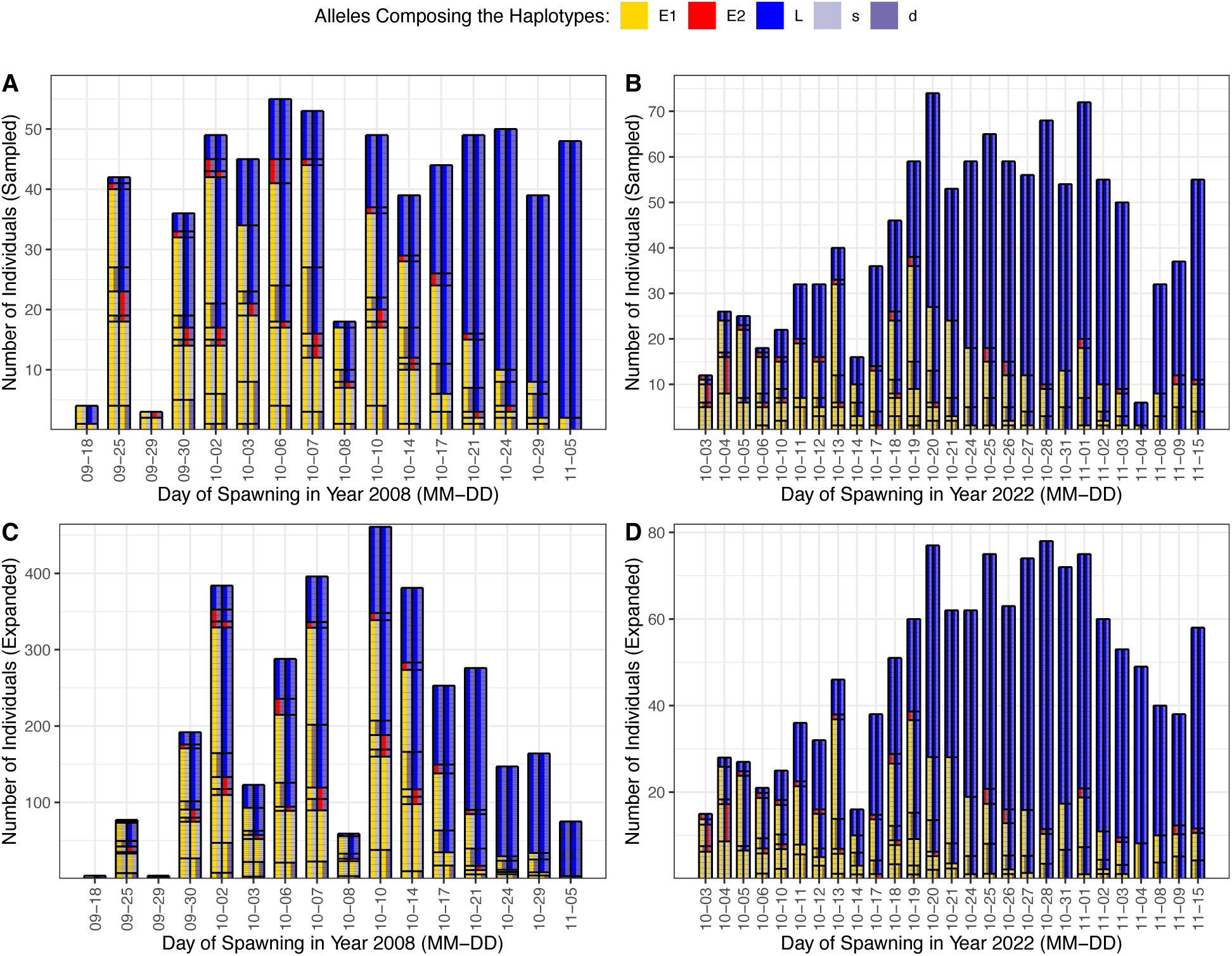
FRH Fall-run program includes many *E* allele homozygotes and heterozygotes. Fall-run spawner RoSA genotypes over time during the FRH fall-run spawning period in 2008 and 2022. Distinct spawning days (listed by date) are arranged on the *x*-axis. In (**A**) and (**B**) the genotype of each sampled individual on a day is represented by a rectangle that is 1 unit (along the y-axis) high. In (**C**) and (**D**), the height of each sampled individual has been expanded by a factor of (Total Fish Spawned on Day / Number of Fish Genotyped on Day), to provide an estimate of the total number of spawners each day of each genotype. Each of these rectangles consists of four squares whose colors denote the alleles carried by the fish. Going from left to right, squares 1 and 3 represent alleles *E1* or *E2* or *L* at the first domain in the *GREB1L*/*ROCK1* region, and squares 2 and 4 represent the *d* or *s* alleles at the second domain. Genotypes have been sorted in the columns so that identical genotypes are adjacent to one another. Thick horizontal lines separate different sets of identical genotypes in the columns and thin horizontal lines separate individual fish. Of note in this figure is the large number of alleles and genotypes associated with early migrating fish found amongst the spawners of the FRH fall-run production as well as the shift over time in the *GREB1L*/*ROCK1* genotypic composition of the spawners and the abundance differences (variable *y*-axis limits) among years.

**Fig. S9:**
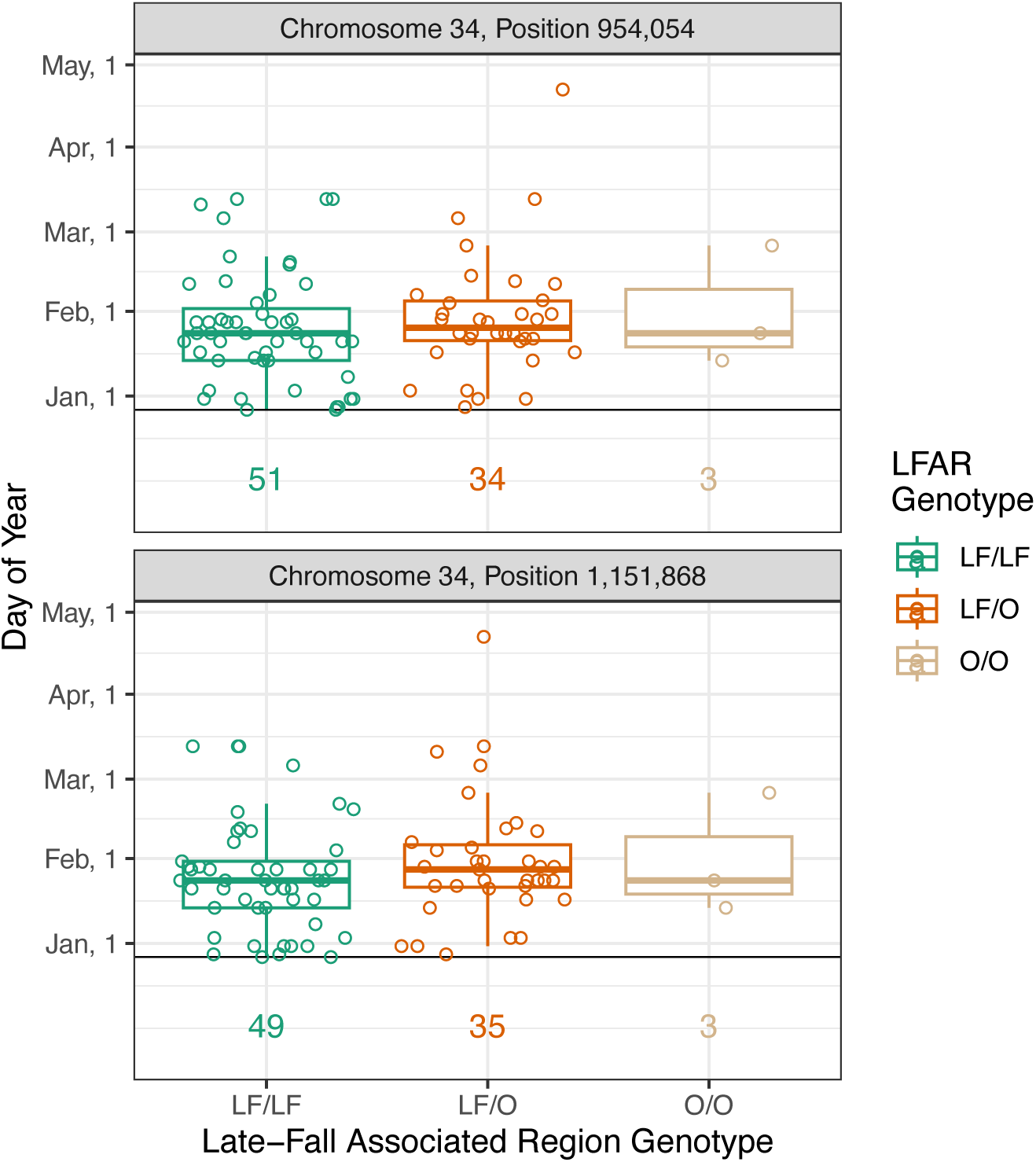
Different genotypes at the late-fall associated region (LFAR) have similar migration timings in the late-fall run. Encounter dates (*y*-axis) of late-fall-run fish at the KFT, grouped by genotypes at two SNPs within the LFAR (*x*-axis). Genotypes are coded according to the allele found at high frequency in the late-fall run (LF) versus the allele found at low frequency in the late-fall run (O). In the boxplots, dark lines are the median of sample points, the hinges are the first and third quartiles and the whiskers extend to the lowest and highest points less than 1.5 times the interquartile range from the lower or upper hinge, respectively. All sample points are plotted under each boxplot. Sampling commences on Dec. 26, as indicated by the thin horizontal line on the plot. SNP positions listed are from the Otsh_v2.0 assembly.

**Tab S1:**
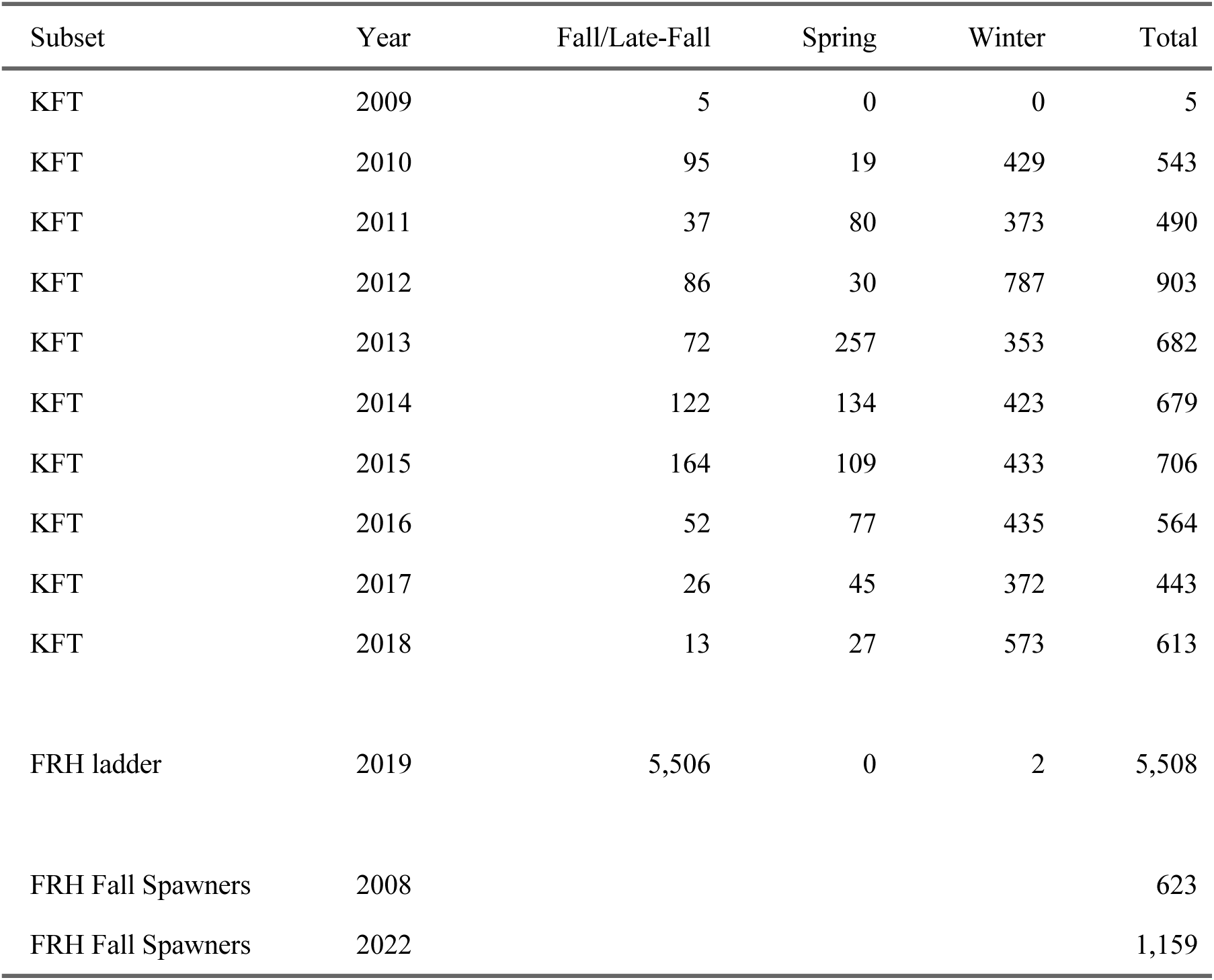
Number of Chinook salmon samples used in analyses of adult migration timing and spawn date. KFT = Keswick Fish Trap and FRH Ladder is the entry ladder to the Feather River Hatchery (FRH). Samples from these locations were used for modeling the relationship between adult migration timing and genetic variation at the *LE1E2* and *sd* domains of the RoSA. FRH Fall Spawners are fish spawned in the fall-run supplementation program at the FRH. Numbers include all fish that passed filters for amount of missing data. GB (Fall/Late-Fall, Spring, Winter), where listed, was inferred from the 125-locus genetic baseline.

**Tab S2:**
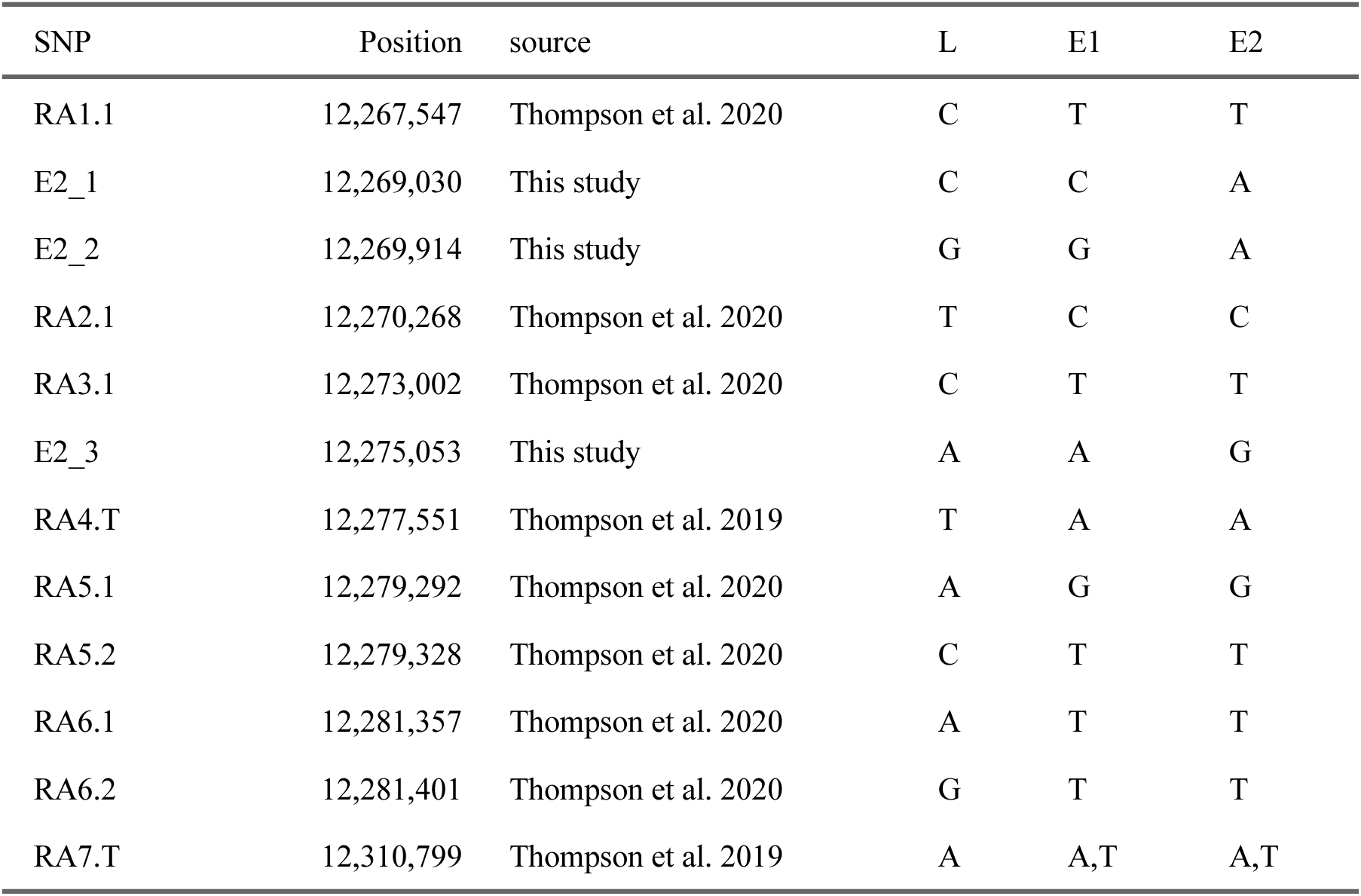
SNP Markers in the RoSA. Genomic coordinates (within Otsh_v1.0, Genbank Accession: GCA_002872995.1), the studies in which they were first described, and the bases found on each of the three haplotypes (*L, E1, E2*) of the 12 SNP markers used to categorize haplotypes in the RoSA. The *L* haplotype always carries an A base at RA7.T; however the *E1* and *E2* alleles may carry either either an A or a T base at RA7.T. At locus RA7.T, the A allele tags the intergenic duplications (*d*) just downstream, while the T allele tags the haplotype that lacks the duplications (*s*).

**Tab S3:**
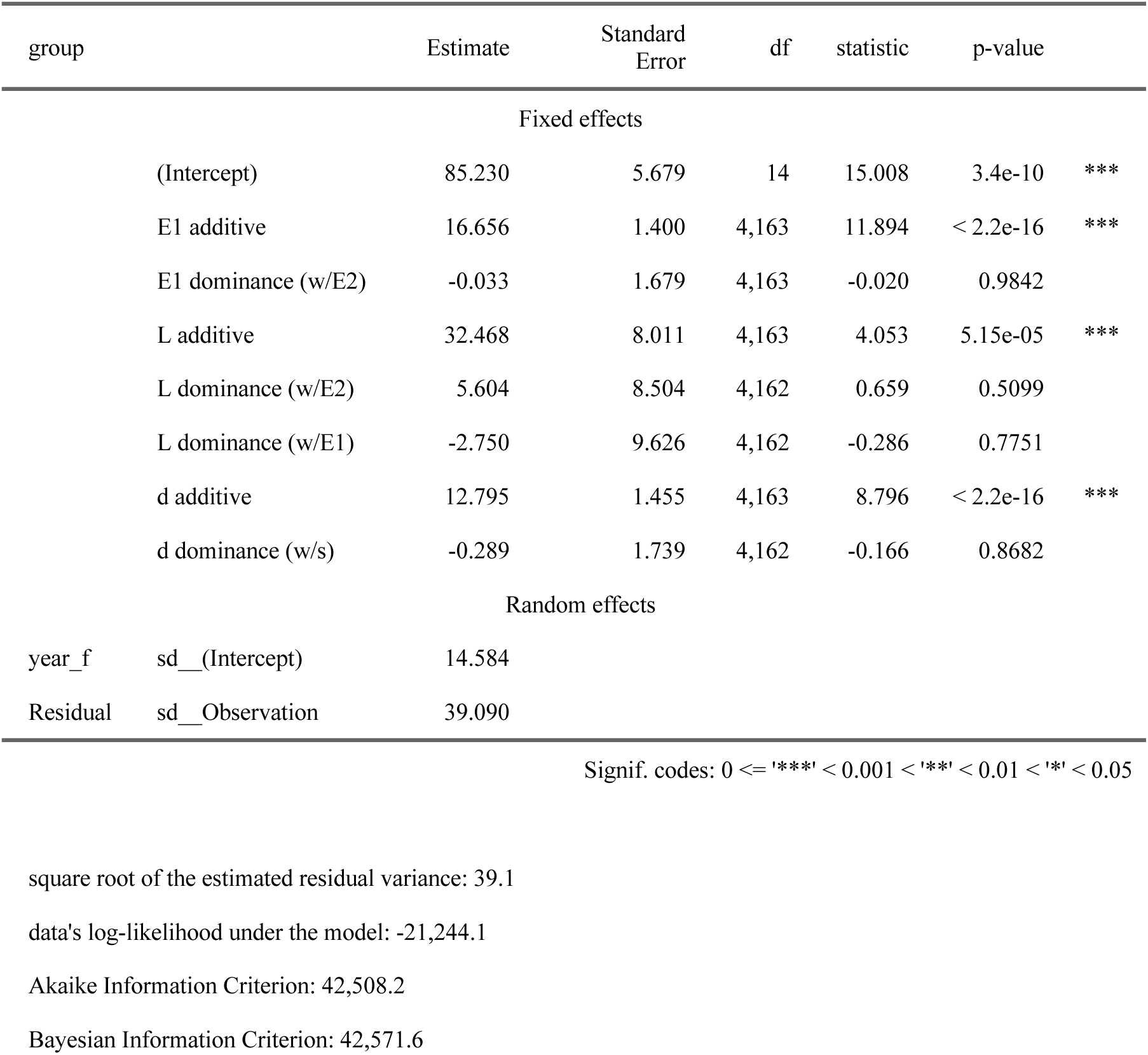
Additive and dominance effects of RoSA alleles in winter-run at KFT. Results of a linear mixed model with year as a random effect and the effect of alleles at the *E1E2L* and *sd* domains captured with a NOIA model. Reference alleles were *E2* at the *E1E2L* domain and *s* at the *sd* domain. Additive effects are all significant. Dominance effects are listed as (w/X) when occurring in heterozygous form with allele X. All dominance effects are non-significant.

**Tab S4:**
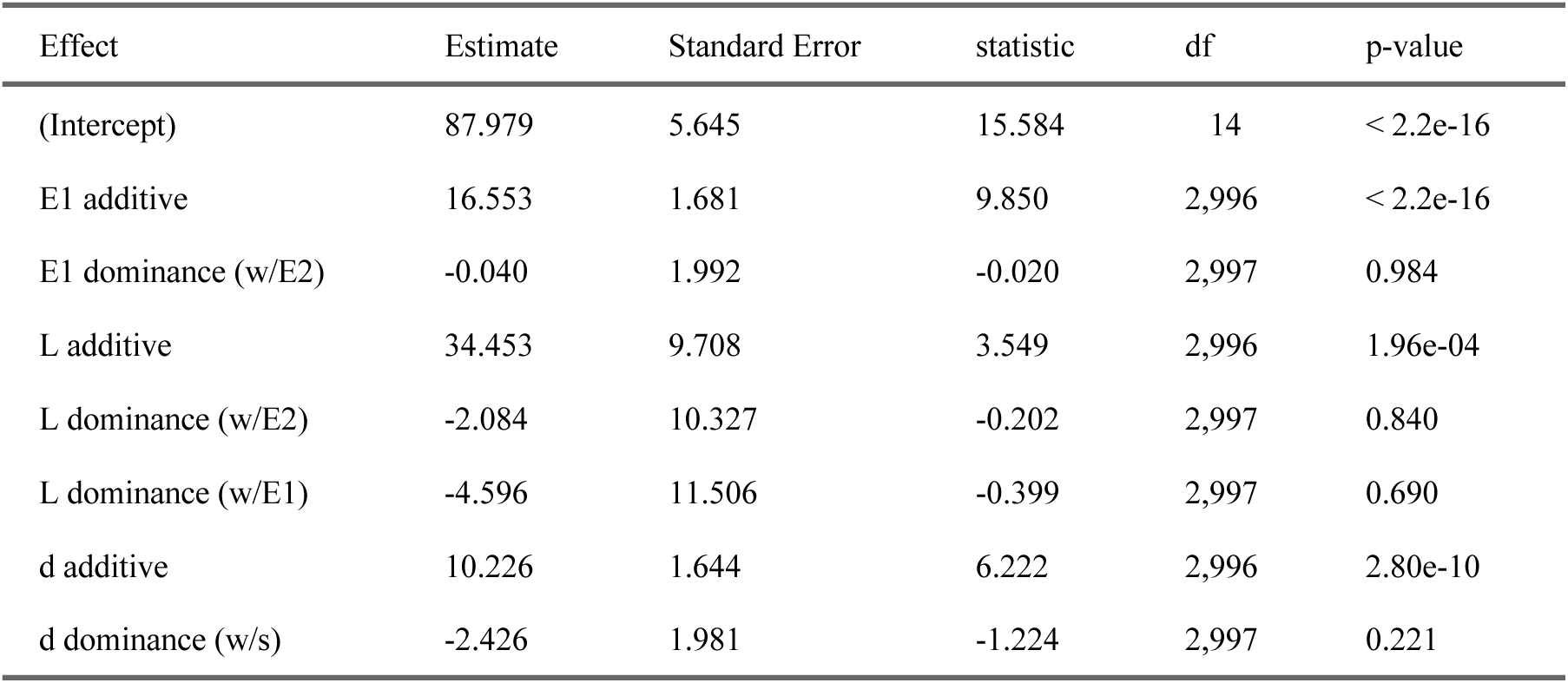
Fixed effects of RoSA alleles in winter-run at KFT accounting for relatedness. Results of a linear mixed model with year and a genomic relationship matrix (calculated from 125 microhaplotypes) as random effects and the effect of alleles at the *E1E2L* and *sd* domains captured with a NOIA model. Reference alleles were *E2* at the *E1E2L* domain and *s* at the *sd* domain. All additive effects are significant. Dominance effects are listed as (w/X) when occurring in heterozygous form with allele X. All dominance effects are non-significant.

**Tab S5:**
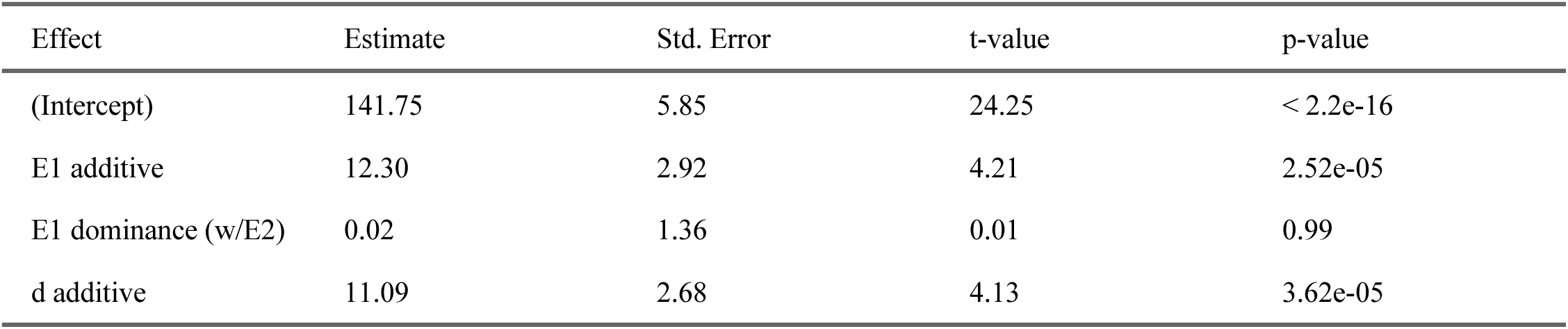
Additive effects at the RoSA estimated from arrival of adult Chinook salmon at the FRH ladder. Arrival time at the ladder was modeled using the NOIA (see Methods) model with only additive effects within a truncated regression framework.

**Tab S6:**
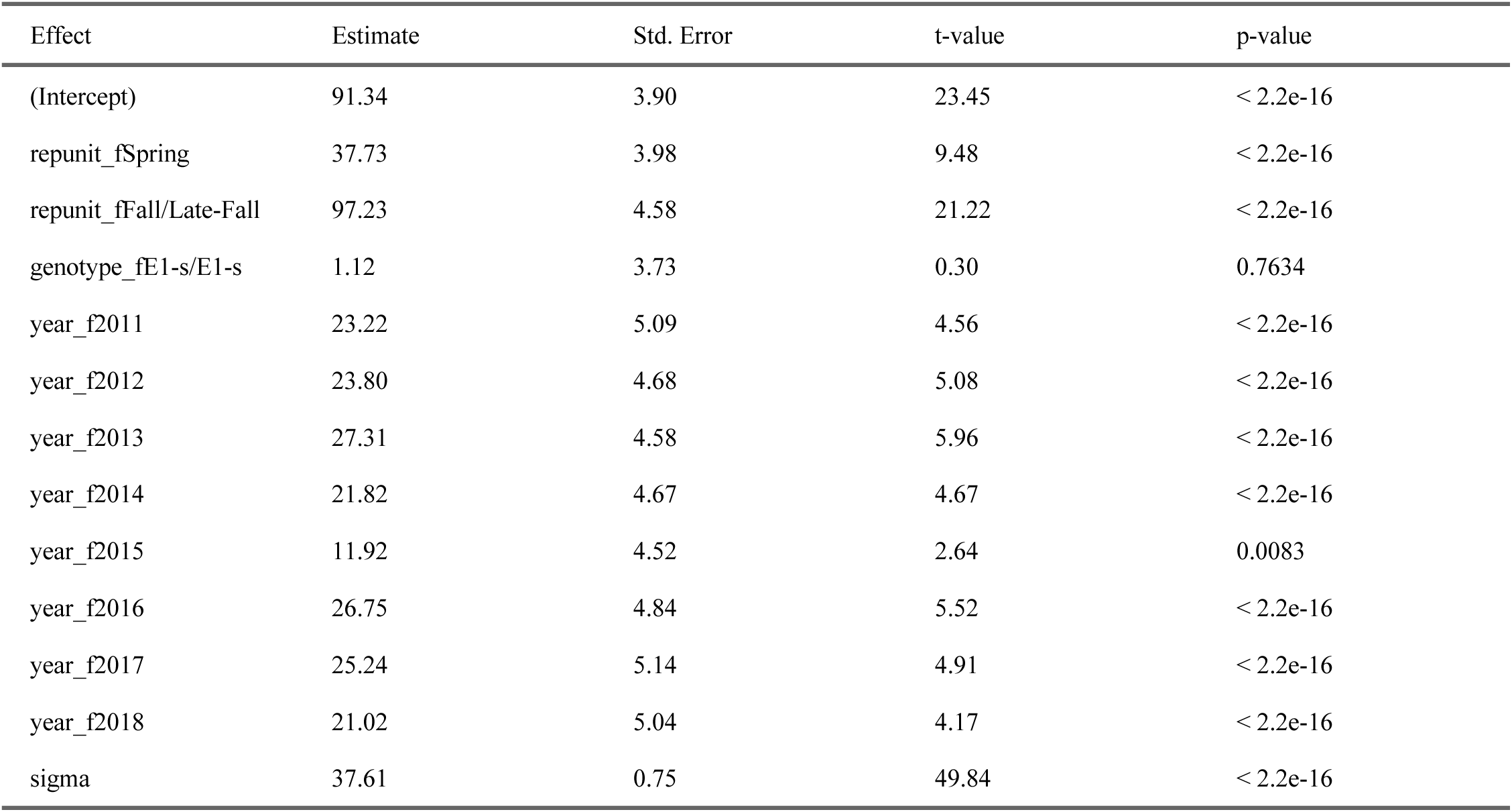
Additive effects of genomic background inferred from arrival times of adult Chinook salmon at KFT. All individuals had either genotype *E1/E2,d/s* or *E1-s/E1-s*. Effects estimated using a truncated regression.

**Tab S7:** GRR Genotype Frequencies in the FRH Fall-run Program Spawners. Counts and relative frequencies of numbers of genotypes at the *LE1E2* and *ds* domains. Genotypes have been sorted first descending by the number of *L* alleles, and then descending on relative frequency. Counts and frequencies have been expanded to account for varying sampling intensity on different days.

| Genotype | 2008 Count | 2022 Count | 2008 Freq | 2022 Freq |
| --- | --- | --- | --- | --- |
| L-d/L-d | 1014.5 | 818.7 | 0.309 | 0.615 |
| E1-s/L-d | 906.2 | 268.9 | 0.276 | 0.202 |
| E1-d/L-d | 322.5 | 103.4 | 0.098 | 0.078 |
| E2-d/L-d | 87.6 | 26.3 | 0.027 | 0.020 |
| E1-s/E1-s | 565.3 | 61.8 | 0.172 | 0.046 |
| E1-s/E1-d | 219.3 | 27.3 | 0.067 | 0.020 |
| E1-s/E2-d | 97.8 | 19.1 | 0.030 | 0.014 |
| E1-d/E2-d | 52.4 | 2.4 | 0.016 | 0.002 |
| E1-d/E1-d | 10.6 | 2.1 | 0.003 | 0.002 |
| E2-d/E2-d | 7.8 | 1.0 | 0.002 | 0.001 |

**Tab S8:**
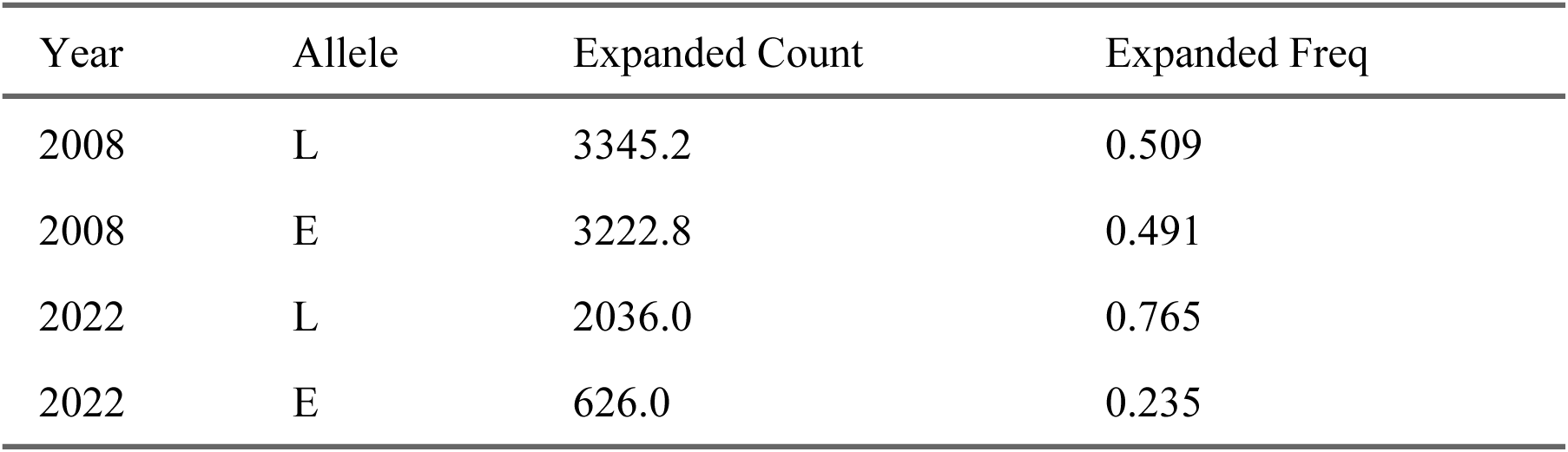
GRR Allele Frequencies in the FRH Fall-run Program Spawners. Counts and relative frequencies of numbers of alleles within spawners that are late-(*L*) or early-(*E = E1 or E2*) associated haplotypes. Values were expanded to account for variable sampling intensity on different days.

## References

1. H. Dingle, Animal migration: Is there a common migratory syndrome? Journal of Ornithology 147, 212–220 (2006).

2. A. J. van Noordwijk, F. Pulido, B. Helm, T. Coppack, J. Delingat, H. Dingle, A. Hedenström, H. van der Jeugd, C. Marchetti, A. Nilsson, J. Pérez-Tris, A framework for the study of genetic variation in migratory behaviour. Journal of Ornithology 147, 221–233 (2006).

3. M. Liedvogel, S. Åkesson, S. Bensch, The genetics of migration on the move. Trends in Ecology & Evolution 26, 561–569 (2011).

4. D. E. Pearse, N. J. Barson, T. Nome, G. Gao, M. A. Campbell, A. Abadía-Cardoso, E. C. Anderson, D. E. Rundio, T. H. Williams, K. A. Naish, T. Moen, S. Liu, M. Kent, M. Moser, D. R. Minkley, E. B. Rondeau, M. S. O. Brieuc, S. R. Sandve, M. R. Miller, L. Cedillo, K. Baruch, A. G. Hernandez, G. Ben-Zvi, D. Shem-Tov, O. Barad, K. Kuzishchin, J. C. Garza, S. T. Lindley, B. F. Koop, G. H. Thorgaard, Y. Palti, S. Lien, Sex-dependent dominance maintains migration supergene in rainbow trout. Nature Ecology & Evolution 3, 1731–1742 (2019).

5. T. Kess, P. Bentzen, S. J. Lehnert, E. V. A. Sylvester, S. Lien, M. P. Kent, M. Sinclair-Waters, C. J. Morris, P. Regular, R. Fairweather, I. R. Bradbury, A migration-associated supergene reveals loss of biocomplexity in Atlantic cod. Science Advances 5, eaav2461 (2019).

6. J. E. Hess, J. S. Zendt, A. R. Matala, S. R. Narum, Genetic basis of adult migration timing in anadromous steelhead discovered through multivariate association testing. Proceedings of the Royal Society B: Biological Sciences 283, 20153064 (2016).

7. D. J. Prince, S. M. O’Rourke, T. Q. Thompson, O. A. Ali, H. S. Lyman, I. K. Saglam, T. J. Hotaling, A. P. Spidle, M. R. Miller, The evolutionary basis of premature migration in Pacific salmon highlights the utility of genomics for informing conservation. Science Advances 3, e1603198 (2017).

8. N. F. Thompson, E. C. Anderson, A. J. Clemento, M. A. Campbell, D. E. Pearse, J. W. Hearsey, A. P. Kinziger, J. C. Garza, A complex phenotype in salmon controlled by a simple change in migratory timing. Science 370, 609–613 (2020).

9. K. Sokolovskis, M. Lundberg, S. Åkesson, M. Willemoes, T. Zhao, V. Caballero-Lopez, S. Bensch, Migration direction in a songbird explained by two loci. Nature Communications 14, 165 (2023).

10. T. P. Quinn, The Behavior and Ecology of Pacific Salmon and Trout (University of Washington Press, 2018).

11. R. L. Horn, S. R. Narum, Genomic variation across Chinook salmon populations reveals effects of a duplication on migration alleles and supports fine scale structure. Molecular Ecology 32, 2818–2834 (2023).

12. A. Abadía-Cardoso, E. C. Anderson, D. E. Pearse, J. C. Garza, Large-scale parentage analysis reveals reproductive patterns and heritability of spawn timing in a hatchery population of steelhead (*Oncorhynchus mykiss*). Molecular Ecology 22, 4733–4746 (2013).

13. A. K. Beulke, A. Abadía-Cardoso, D. E. Pearse, L. C. Goetz, N. F. Thompson, E. C. Anderson, J. C. Garza, Distinct patterns of inheritance shape life-history traits in steelhead trout. Molecular Ecology 32, 6896–6912 (2023).

14. R. M. Yoshiyama, F. W. Fisher, P. B. Moyle, Historical abundance and decline of Chinook salmon in the Central Valley region of California. North American Journal of Fisheries Management 18, 487–521 (1998).

15. E. C. Anderson, A. J. Clemento, M. A. Campell, D. E. Pearse, A. K. Beulke, C. Columbus, E. Campbell, N. F. Thompson, J. C. Garza, A multipurpose microhaplotype panel for genetic analysis of California Chinook salmon. Evolutionary Applications 18, e70110 (2025).

16. P. W. Hedrick, D. Hedgecock, Effective population size in winter-run Chinook salmon. Conservation Biology 8, 890–892 (1994).

17. A. J. Clemento, E. D. Crandall, J. C. Garza, E. C. Anderson, Evaluation of a single nucleotide polymorphism baseline for genetic stock identification of Chinook Salmon (*Oncorhynchus tshawytscha*) in the California Current large marine ecosystem. Fishery Bulletin 112 (2014).

18. L. E. Kruuk, Estimating genetic parameters in natural populations using the “animal model.” Philosophical Transactions of the Royal Society of London. Series B: Biological Sciences 359, 873–890 (2004).

19. A. J. Wilson, D. Réale, M. N. Clements, M. M. Morrissey, E. Postma, C. A. Walling, L. E. Kruuk, D. H. Nussey, An ecologist’s guide to the animal model. Journal of Animal Ecology 79, 13–26 (2010).

20. C. Bérénos, P. A. Ellis, J. G. Pilkington, J. M. Pemberton, Estimating quantitative genetic parameters in wild populations: A comparison of pedigree and genomic approaches. Molecular Ecology 23, 3434–3451 (2014).

21. T. A. Manolio, F. S. Collins, N. J. Cox, D. B. Goldstein, L. A. Hindorff, D. J. Hunter, M. I. McCarthy, E. M. Ramos, L. R. Cardon, A. Chakravarti, J. H. Cho, A. E. Guttmacher, A. Kong, L. Kruglyak, E. Mardis, C. N. Rotimi, M. Slatkin, D. Valle, A. S. Whittemore, M. Boehnke, A. G. Clark, E. E. Eichler, G. Gibson, J. L. Haines, T. F. C. Mackay, S. A. McCarroll, P. M. Visscher, Finding the missing heritability of complex diseases. Nature 461, 747–753 (2009).

22. A. P. Kinziger, M. Hellmair, D. G. Hankin, J. C. Garza, Contemporary population structure in Klamath River basin Chinook salmon revealed by analysis of microsatellite genetic data. Transactions of the American Fisheries Society 142, 1347–1357 (2013).

23. G. Perez, G. P. Barber, A. Benet-Pages, J. Casper, H. Clawson, M. Diekhans, C. Fischer, J. N. Gonzalez, A. S. Hinrichs, C. M. Lee, L. R. Nassar, B. J. Raney, M. L. Speir, M. J. van Baren, C. J. Vaske, D. Haussler, W. J. Kent, M. Haeussler, The UCSC genome browser database: 2025 update. Nucleic Acids Research 53, D1243–D1249 (2024).

24. M. Gardiner-Garden, M. Frommer, CpG islands in vertebrate genomes. Journal of molecular biology 196, 261–282 (1987).

25. T. Q. Thompson, M. R. Bellinger, S. M. O’Rourke, D. J. Prince, A. E. Stevenson, A. T. Rodrigues, M. R. Sloat, C. F. Speller, D. Y. Yang, V. L. Butler, M. A. Banks, M. R. Miller, Anthropogenic habitat alteration leads to rapid loss of adaptive variation and restoration potential in wild salmon populations. Proceedings of the National Academy of Sciences 116, 177–186 (2019).

26. D. A. Vogel, M. K. R, “Guide to Upper Sacramento River Chinook Salmon Life History. Prepared for the U.S. Bureau of Reclamation, Central Valley Project.” (CH2M, 1991).

27. G. B. Milner, D. J. Teel, F. M. Utter, G. A. Winans, A genetic method of stock identification in mixed populations of Pacific salmon, Oncorhynchus spp. Marine Fisheries Review 47, 1–8 (1985).

28. D. Paetkau, R. Slade, M. Burden, A. Estoup, Genetic assignment methods for the direct, real-time estimation of migration rate: A simulation-based exploration of accuracy and power. Molecular Ecology 13, 55–65 (2004).

29. D. S. Baetscher, A. J. Clemento, T. C. Ng, E. C. Anderson, J. C. Garza, Microhaplotypes provide increased power from short-read DNA sequences for relationship inference. Molecular Ecology Resources 18, 296–305 (2018).

30. B. M. Moran, E. C. Anderson, Bayesian inference from the conditional genetic stock identification model. Canadian Journal of Fisheries and Aquatic Sciences 76, 551–560 (2019).

31. T. S. Korneliussen, A. Albrechtsen, R. Nielsen, ANGSD: Analysis of next generation sequencing data. BMC Bioinformatics 15, 356 (2014).

32. M. S. Rasmussen, G. Garcia-Erill, T. S. Korneliussen, C. Wiuf, A. Albrechtsen, Estimation of site frequency spectra from low-coverage sequencing data using stochastic EM reduces overfitting, runtime, and memory usage. Genetics 222, iyac148 (2022).

33. H. Li, B. Handsaker, A. Wysoker, T. Fennell, J. Ruan, N. Homer, G. Marth, G. Abecasis, R. Durbin, 1000. G. P. D. P. Subgroup, The sequence alignment/map format and SAMtools. Bioinformatics 25, 2078–2079 (2009).

34. A. G. Clark, Inference of haplotypes from PCR-amplified samples of diploid populations. Molecular Biology and Evolution 7, 111–122 (1990).

35. H. Wickham, Ggplot2: Elegant Graphics for Data Analysis (Springer-Verlag New York, 2016; https://ggplot2.tidyverse.org).

36. J. M. Álvarez-Castro, Ö. Carlborg, A unified model for functional and statistical epistasis and its application in quantitative trait loci analysis. Genetics 176, 1151– 1167 (2007).

37. R.-C. Yang, J. M. Álvarez-Castro, Functional and statistical genetic effects with multiple alleles. Current Topics in Genetics 3, 49–62 (2008).

38. Y. Croissant, A. Zeileis, Truncreg: Truncated Gaussian Regression Models (2018; https://CRAN.R-project.org/package=truncreg).

39. C.-P. Giovanny, Genome assisted prediction of quantitative traits using the R package sommer. PLoS ONE 11, 1–15 (2016).

40. P. M. VanRaden, Efficient methods to compute genomic predictions. Journal of Dairy Science 91, 4414–4423 (2008).

41. T. H. Meuwissen, J. Odegard, I. Andersen-Ranberg, E. Grindflek, On the distance of genetic relationships and the accuracy of genomic prediction in pig breeding. Genetics Selection Evolution 46, 1–8 (2014).

